# Integrative ensemble modelling of cetuximab sensitivity in colorectal cancer PDXs

**DOI:** 10.1101/2023.01.24.525314

**Authors:** Umberto Perron, Elena Grassi, Aikaterini Chatzipli, Marco Viviani, Emre Karakoc, Lucia Trastulla, Claudio Isella, Eugenia R Zanella, Hagen Klett, Ivan Molineris, Julia Schueler, Manel Esteller, Enzo Medico, Nathalie Conte, Ultan McDermott, Livio Trusolino, Andrea Bertotti, Francesco Iorio

**Affiliations:** Human Technopole, 20157, Milano, Italy; Candiolo Cancer Institute FPO IRCCS, 10060 Candiolo, Torino, Italy; Department of Oncology, University of Torino, 10060 Candiolo, Torino, Italy; Wellcome Sanger Institute, Cambridge, UK; Charles River Germany GmbH, Freiburg, Germany; European Molecular Biology Laboratory - European Bioinformatics Institute, Cambridge, United Kingdom; Department of Life Sciences and Systems Biology, University of Torino, Torino, Italy; IIGM - Italian Institute for Genomic Medicine, c/o IRCCS, Candiolo, Italy; Josep Carreras Leukemia Research Institute (IJC), Badalona, Barcelona, Catalonia, Spain; Centro de Investigacion Biomedica en Red Cancer (CIBERONC), 28029 Madrid, Spain; Institucio Catalana de Recerca i Estudis Avançats (ICREA), Barcelona, Catalonia, Spain; Physiological Sciences Department, School of Medicine and Health Sciences, University of Barcelona (UB), Barcelona, Catalonia, Spain

**Author notes:** co-first authors (Umberto Perron, Elena Grassi, Aikaterini Chatzipli).

**Keywords:** *colorectal-cancer*, *PDXs*, *drug-response*, *predictive-model*, *EGFR*

## Abstract

Patient-derived xenografts (PDXs) are tumour fragments engrafted into mice for preclinical studies. PDXs offer clear advantages over simpler *in vitro* cancer models - such as cancer cell lines (CCLs) and organoids - in terms of structural complexity, heterogeneity, and stromal interactions. We characterised 231 colorectal cancer PDXs at the genomic, transcriptomic, and epigenetic level and measured their response to cetuximab, an EGFR inhibitor in clinical use for metastatic colorectal cancer. After assessing PDXs’ quality, stability, and molecular concordance with publicly available patient cohorts, we trained, interpreted, and validated an integrated ensemble classifier (CeSta) which takes in input the PDXs’ *multi-omic* characterisation and predicts their sensitivity to cetuximab treatment (AUROC > 0.9). Our study shows that large PDX collections can be used to train accurate, interpretable models of drug sensitivity, which 1) better recapitulate patient-derived therapeutic biomarkers than other models trained on CCL data, 2) can be robustly validated across independent PDX cohorts, and 3) can be used for the development of novel therapeutic biomarkers.

## Introduction

Colorectal cancer (CRC) is a heterogeneous disease with distinctly variable molecular features and responses to therapy. It is among the most prevalent causes of cancer mortality worldwide, with more than 1.85 million cases and 850,000 annual deaths globally^1^. Around 20% of newly diagnosed CRC patients have metastatic disease (mCRC) at presentation, with 25% later developing metastases^2–4^. In recent years, several clinical trials^5–7^ have suggested that genome-based treatment selection leads to more patients deriving therapeutic benefits, with fewer exposed to ineffective therapies, and most mCRC patients experience a median survival exceeding 30 months when regimens including genotype-informed treatments are used^8^. Specifically, ∼50% of mCRC patients have KRAS-NRAS-BRAF wild-type (triple negative) tumours and are routinely treated with cetuximab and panitumumab, monoclonal antibody inhibitors of the epithelial growth factor receptor EGFR in combination with chemotherapy as an alternative to surgery. This protocol extends median survival by 2 to 4 months, compared with chemotherapy alone^1^. Unfortunately, the overall metastatic CRC clinical trial success rate remains low: 32% of combined phase II and phase III clinical trials between 2013 and 2015 failed, up from 23% in 2010^9^. This highlights the need for novel and more robustly predictive markers of drug response for CRC patients.

Biomarkers of response to cetuximab and cetuximab plus chemotherapy, such as the triple negative signature mentioned above, have been derived from clinical and molecular analysis of patients and patient-derived experimental models of CRC, including immortalised cancer cell lines, organoids, and patient-derived xenografts (PDX). However, several other systematic therapeutic biomarkers discovery efforts conducted using *in vitro* models have confirmed limited clinical translatability^9–11^. This is primarily due to the intrinsic limitations of such models, encompassing genetic, epigenetic, and transcriptomic changes resulting from their selective adaptation to artificial culture conditions^12, 13^. Furthermore, cancer cell lines do not maintain the complex heterogeneity of the tumour of derivation; they often lose or gain specific subclones and might miss relevant components of the human tumour stromal microenvironment^14, 15^.

Unlike cancer cell lines, PDXs have been shown to offer good retention of tumour complexity, mimicking (at least to a certain extent) stromal interactions. They are relatively easy to screen and characterise. Further, histopathological characterisation has confirmed a high degree of concordance between PDXs and corresponding parental tumours in terms of differentiation, mucus secretion, and stromal composition, as well as maintenance of primary intratumoral clonal heterogeneity^2, 3, 16–18^.

These factors have contributed to PDXs playing a pivotal role in translational cancer research, furthering our understanding of tumour biology and drug response mechanisms in CRC^19, 20^. As a result, extensive multi-institutional efforts (such as EuroPDX^21^) are now ongoing, aiming to establish and characterise extensive collections of PDX models at the molecular and histopathological level to ensure that they recapitulate the broadest possible diversity of clinical cases^22^.

Using data derived from the multi-omics characterisation of CRC PDXs paired with their pharmacological/phenotypic features is a profitable means for training supervised machine learning models to predict drug response in CRC patients. In this case, the extent of training data availability is a critical determinant of the accuracy of a model, especially when considering high-dimensional multi-omics datasets. Machine learning models of drug response trained on large pooled pan-cancer cell line datasets (N = 329) outperform models which only used cell lines (N = 28-68) from a specific tissue^23^. This suggests that, in some cases, data quantity can outweigh data specificity. Kurilov and colleagues have also noted that predicting PDX drug response using models trained on cell line data results in poor performance across 3 out of 4 examined cohorts, except for the erlotinib lung cancer cohort.

In summary, most of the pre-clinical studies of cetuximab response in CRC cohorts performed to date have been characterised by 1) relatively small sample sizes, 2) single platform profiling often aimed at characterising the status of few known CRC driver genes, 3) reliance on biological models which have proved to be suboptimal for translational purposes, or a combination of these factors. These aspects negatively influenced the aforementioned studies’ ability to capture the tumour ecosystem’s complexity and inter-tumour heterogeneity’s impact on drug response, ultimately contributing to the increasingly low success rate of early-stage CRC clinical trials.

Here, we present one of the largest thoroughly characterised CRC PDX collections to date (N = 231), which closely recapitulates gold-standard CRC patient cohorts across 3 ‘omics (genomics, transcriptomics, and methylomics) and results from training an ensemble classifier to predict the response of these models to cetuximab treatment, based on an integrative stacked architecture.

Our model outperforms other state-of-the-art (SOTA) predictive methods and the biomarker of cetuximab response currently used in the clinic, i.e. the KRAS-NRAS-BRAF mutational status, internally and when tested on an independent cohort of CRC PDXs.

Finally, we show that our model’s predictions provide an extent of interpretability, highlighting novel potential biomarkers of cetuximab sensitivity.

## Results

We selected 231 first-pass CRC PDXs (the IRCC-PDX collection), which were fully characterised across multiple omics (encompassing genomics, transcriptomics, methylomics), clinical metadata, and were screened for cetuximab response from a larger cohort of >600 xenografts (**Fig. 1a**). These models were uniquely derived from surgical resections of CRC liver metastases performed at the Candiolo Cancer Institute (Candiolo, Torino, IT), the Mauriziano Umberto I Hospital (Torino, IT), the San Giovanni Battista Hospital (Torino, IT) and the Niguarda Hospital (Milano, IT) between 2008 and 2015.

**Figure 1.**
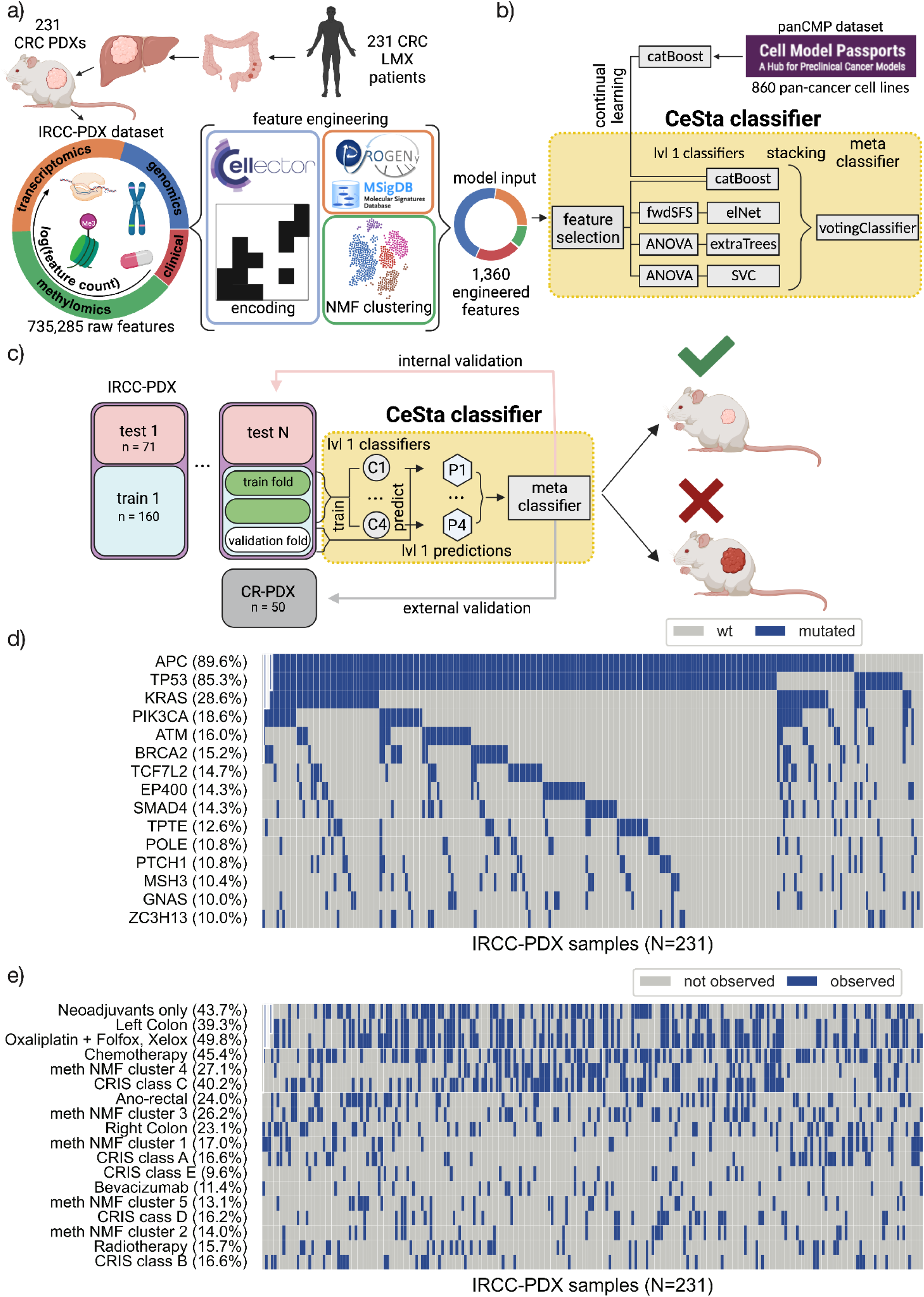
A multi-omic overview of the Colorectal cancer PDX cohort and cetuximab response modelling approach. **a)** Overview of the IRCC-PDX collection, derived from 231 unique CRC liver metastasis resections, characterised at a multi-omic level as well as for cetuximab response, including a schematic of the omic-specific feature engineering steps. **b)** Overview of our CeSta classifier pipeline. A set of input features is selected from the training set (Methods) using univariate tests (Fisher’s exact, MW U-test) and multivariate linear models. These features are passed as input to 3 independent level 1 classifier pipelines (lvl 1 classifiers): forward feature selection plus elastic net classifier, ANOVA feature selection plus extra trees classifier, ANOVA feature selection plus SV classifier. An additional fourth classifier uses a boosting (catBoost) model, which we pre-train using pan-cancer data from the cell model passport repository (CMP), and we then refine using the same IRCC-PDX data as the other lvl 1 classifiers (Methods). Lvl 1 classifier predicted probabilities are stacked and provided as input to a meta classifier which outputs the final binary class label (i.e. cetuximab-responder/-non-responder) corresponding to the lvl 1 prediction with the highest predicted probability (i.e. argmax-based soft voting). **c)** CeSta nested cross-validation approach: 50 train/test split replicates are derived by stratified sampling of the IRCC-PDX collection. CeSta is trained and tuned independently across these 50 data splits. In each of these iterations, the training set is split into three folds, two of which are used (in turn) in three successive rounds, jointly as a “training fold” (green rectangles). In each of these rounds, the level-1-classifiers generate predictions for the remaining subset not used for model fitting, i.e. the “validation fold” (white rectangle). The resulting predictions (“P1” and “P2”, green hexagon) are then stacked and provided to the meta-classifier. After comparing the meta-classifier’s prediction on the validation fold to the corresponding true labels, all first-level classifiers are fit to the entire training set replicate and CeSta performance is evaluated on the test set (pink rectangle, N = 71) of the split under consideration (“internal validation”). Finally, CeSta is trained once over the entire IRCC-PDx dataset and tested (“external validation”) on an independent CR-PDX dataset (grey rectangle, N=50). **d)** Top frequently mutated genes in our IRCC-PDX cohort. **e)** Selection of multi-omic and clinical features across the IRCC-PDX collection, including CRIS expression cluster labels, methylation NMF cluster labels, primary sample anatomical location, and treatment backbone.

The initial “raw” multi-omics characterisation of IRCC-PDX consisted of the methylation status of 700,298 Illumina probes, 33,670 gene transcription levels from RNAseq, 1,272 copy number alteration and driver variant features, and 45 clinical features covering patient demographics, primary tumour characteristics, and previous patient treatment for a total of 735,285 features (**Fig. 1a**).

We performed several omic-specific feature engineering steps (Methods, **Fig. 1a**) before using this data with our integrative classifier (**Fig. 1b-c**). These reduced the dimensionality of the “raw” IRCC-PDX dataset (e.g. non-negative matrix factorisation clustering^24^ of methylation features), introduced feature curation via prior knowledge of gene regulatory pathways (e.g. PROGENy^25^ and MSigDB^26^ gene set analysis), and generated potentially more informative agglomerate features (e.g. CELLector^27^ genomic signatures). Raw and engineered (1,360 features, 231 PDX models) IRCC-PDX datasets are both fully available for download (Data and Code Availability).

### Multi-omic characterisation of our CRC PDX collection

Previous comprehensive genetic characterisations of CRC models have shown that the frequency of common genetic mutations observed in PDXs is similar to that observed in primary tumours^2, 3, 16–18, 28^. Targeted sequencing of 116 genes in our PDX cohort identified 6,426 driver mutations (Methods), with APC (90%), TP53 (85%), KRAS (29%), PIK3CA (19%), and ATM (16%) being the most frequently affected genes (**Fig 1d**, **S1**). In our PDX collection, mutational frequencies for KRAS and BRAF were lower than those reported for large CRC patient cohorts such as TCGA COAD/READ (https://www.cancer.gov/tcga) and MSK IMPACT^29^ (https://www.mskcc.org/msk-impact). In the case of KRAS, this is due to a pre-hoc enrichment of KRAS wild-type models for subsequent treatment with cetuximab (as KRAS mutant models were assumed to be cetuximab resistant a priori). In the case of BRAF, the lower frequency is ascribable to the fact that our PDXs were derived from metastatic samples. BRAF mutant tumours are frequently characterised by microsatellite instability (MSI). Because MSI CRCs have a better prognosis and rarely progress to metastasis^30^, they are under-represented in our dataset. Indeed, after removing MSI samples, the frequency of BRAF mutated tumours in TCGA is reduced to 5.3%, which is comparable to that detected in our collection.

Aside from these exceptions, our IRCC-PDX mutational landscape closely matched that of the previous CRC patient cohorts (Spearman correlation coefficient 0.51 and 0.625 for TCGA and MSK, respectively; **Fig. S2**) and recapitulated known top frequently mutated CRC driver genes^31, 32^.

To further control our PDX models’ ability to recapitulate characteristics of their tumour sample of origin, we investigated PDX mutational profile stability for a subset of more extended PDX lineages (i.e. those where targeted sequencing data was available beyond the first-passage; **Fig. S3**). We observed a strong agreement between all models belonging to a given lineage, regardless of their distance from their sample of origin in terms of passages, with few exceptions attributable to sequencing errors or clonal expansion (**Fig. S4**).

Copy number (CN) alterations, derived from the same 116 genes in the targeted sequencing panel (Methods), affected some known CRC drivers, including EGFR and SMAD4, and showed a strong positive correlation (Spearman r = 0.87 and 0.93, respectively, for copy number losses and gains) with CN alteration frequencies observed in TCGA COAD/READ samples (**Fig. S5** and **S6**).

As described above, we also assessed CN profile stability along PDX lineages which extend beyond the first passage. We observed solid intra-lineage CN consistency overall (median log2R Pearson coefficient 0.927, **Fig. S7**) and at the gene level (94% of driver genes are CN-stable within lineages, **Fig. S8**), in line with previous reports^33^.

We characterised our PDX collection’s transcriptional landscape using two approaches to classify samples into subtypes: CMS^34, 35^ and CRIS^4^. Results from these analyses were broadly consistent with TCGA COAD/READ and other colorectal cancer datasets where expression data is available (**Fig. 1e, S9**).

To concisely represent our PDXs’ epigenomics profiles, we grouped samples into five different nonnegative matrix factorisation^24^ based clusters (Methods). We observed that the samples belonging to one of these groups (cluster 1) were remarkably more hypermethylated over all measured CpG islands (median beta methylation level = 0.81, Kruskal-Wallis test p-value < 2.2e-16, **Fig. S10**). Consistent with our cluster definition, we also found cluster 1 to be highly enriched for the CpG island methylator phenotype (CIMP^36^) in 130 out of 146 PDXs (**Fig. S10**). This heterogeneity of PDX methylation profiles resembled that observed in CRC patients, even though the percentage of IRCC-PDX samples classified as CIMP was slightly lower than that reported in TCGA COAD/READ (44% vs 58%, **Fig. S11**). This is expected considering the low prevalence of MSI tumours - which are typically enriched for CIMP cases - within metastatic CRC cohorts such as ours^29^.

Overall, our multi-omic overview of the PDX collection indicates that IRCC-PDX closely recapitulates the genomics, transcriptomics, and methylomics landscape of gold-standard human colorectal cancer cohorts, such as TCGA COAD/READ and MSK-IMPACT.

### Exploration of established biomarkers of cetuximab sensitivity

Around half of mCRC patients have KRAS-NRAS-BRAF wild-type (triple negative) tumours and routinely receive anti-EGFR treatment with cetuximab or panitumumab in combination with chemotherapy as an alternative to surgery. This results in a median survival extension of 2 to 4 months, compared with chemotherapy alone^1^. Retrospective analysis of triple-negative patients from the CRYSTAL and FIRE3 trials has also highlighted that patients with left-sided tumours treated with anti-EGFR antibodies had better survival and treatment response than patients with right-sided tumours^37^.

Treatment intervention in our PDXs (Methods) closely matched that of cetuximab human trials such as PEAK^7, 38^ and FIRE3^5^ as well as current clinical best practices^39, 40^; in line with the clinical definition of ‘disease control’, which denotes clinical benefit, we categorised as ‘responders’ those cases in which cetuximab induced tumour shrinkage (objective response, OR, more than 50% tumour volume reduction compared with baseline tumour volumes) or stable disease (SD, less than 50% tumour shrinkage and less than 35% increase in tumour volume^2^).

Across our IRCC-PDX collection (N = 231), KRAS mutations were much more frequently observed in PDXs with a cetuximab non-responder phenotype (Fisher’s exact test’s *p-value* (FET p) = 4×10^−6^, percent lift: −0.781, **Fig 2a, Tab S1**, Methods). NRAS (FET p = 0.001, percent lift: −0.926) and BRAF (FET p = 0.035, percent lift: −0.702) mutations were noticeably more likely to occur in non-responder PDXs, though only 16 and 13 mutant PDXs were observed across IRCC-PDXs, respectively. However, overall mutational and CN alteration burden, defined as the total number of events per PDX and intended as coarse-grained proxies for tumour progression and genomic stability, did not appear to strongly correlate with cetuximab sensitivity (**Fig 2b-c, Tab S1**).

**Figure 2.**
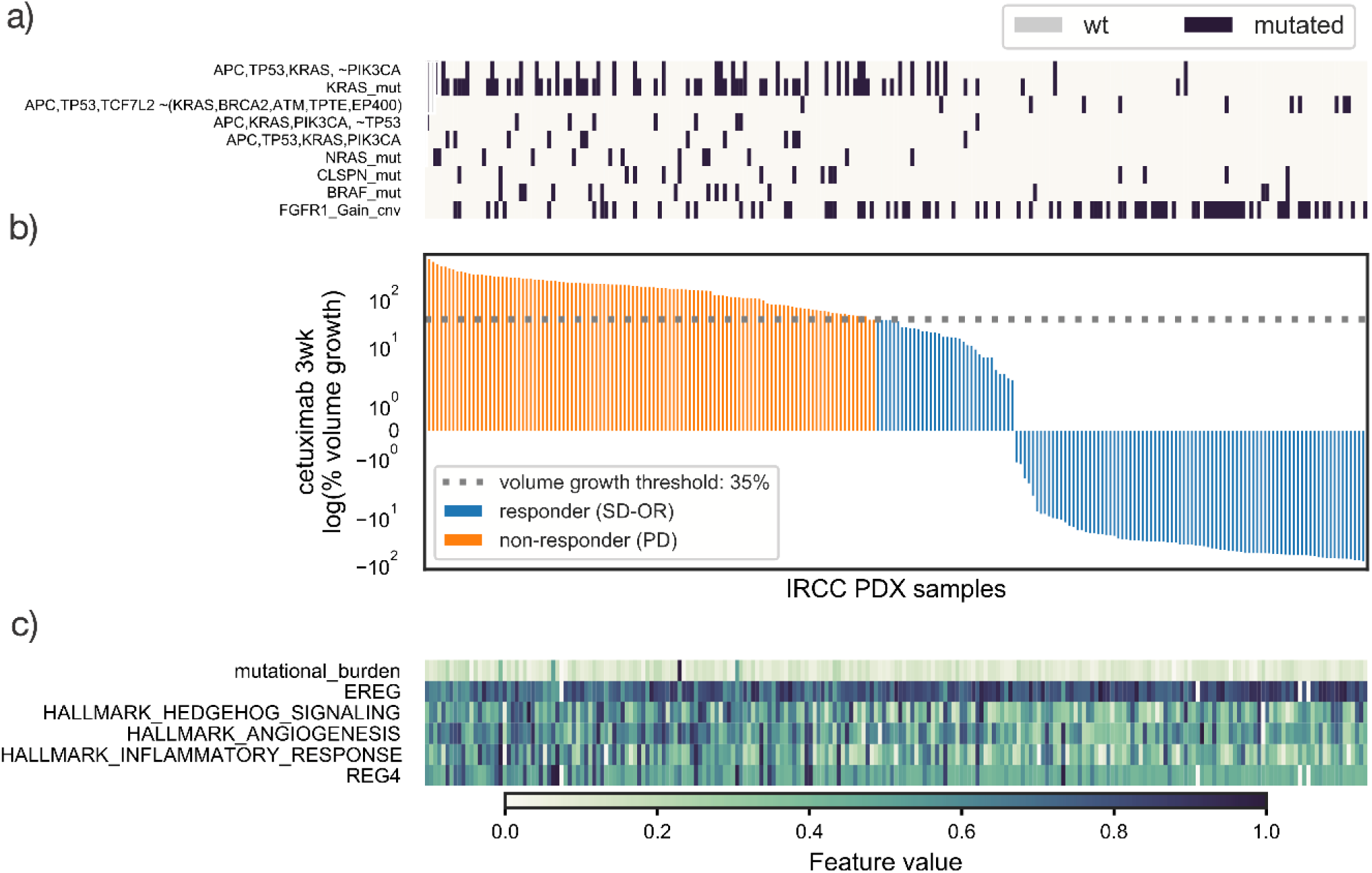
Overview of cetuximab response and biomarker candidates. **a)** Mutation patterns of CRC driver genes and mutational signature features among those with the most significant impact on CeSta predictions (**Fig 4a**) of **b)** cetuximab non-responders (‘PD’, volume growth > 35%, in orange) and responders (‘SD-OR’, volume growth <= 35%, in blue). Similarly, **c)** selection of continuous features which best differentiate between PD and SD-OR PDX models.

Finally, a right-sided localisation of the original tumour showed a moderate association with a non-responder phenotype (FET p = 0.017, percent lift: −0.470).

As previously mentioned, the KRAS-NRAS-BRAF triple negative signature is widely recognised as the best-established biomarker of cetuximab sensitivity (FET p = 5×10^−15^, percent lift: 1.468), being used both as a clinical discriminant for treatment and as an entry criterion for anti-EGFR trials. These observations thus indicate that our IRCC-PDX collection recapitulates the best available marker of cetuximab sensitivity in patients.

### A stacked classifier modelling cetuximab sensitivity

We tackled the task of predicting whether a CRC PDX responds to cetuximab treatment in terms of tumour volume shrinkage^2^ by leveraging its multi-omic characterisation and formulating this challenge as a binary classification problem. This mirrors a clinician’s choice regarding whether or not a patient might benefit from cetuximab treatment.

We selected and integrated multi-omic features into a stacked classifier pipeline^41^: the cetuximab Stacked classifier (CeSta, **Fig 1b**). Stacking is a supervised ensemble learning technique which combines multiple weak classification models (level 1 classifiers, lvl1) using a meta-classifier. This architecture improves upon individual classifiers’ performance. It is well suited for a classification task such as ours, which is based on tabular data with relatively few examples (231) and a much larger number of features (1,360): a scenario where more complex models and deep neural networks fare poorly^42, 43^. A similar architecture has been successfully used to predict drug response in breast cancer patients from the multi-omic characterisation of their tumours^44^.

Our CeSta pipeline implements a late integration approach to prevent high-dimensional ‘omics (transcriptomics, methylomics) from overwhelming those with fewer features (typically genomics) by dominating the feature selection phase (**Fig 1b**). We used a nested cross-validation approach for model tuning, training, and validation, based on generating 50 train/test split replicates of our IRCC-PDX dataset (with 160 and 71 PDXs, respectively, for the training set and test set) assembled via stratified sampling (**Fig 1c**). On each of these 50 training sets, our classifier pipeline performed a custom single omic feature selection step which reduced the initial input of 1,360 engineered features (**Fig. 1a**) to a smaller subset, with the size of the latter being amongst the hyperparameters tuned independently, across data splits (**Fig. 1b, S16,** Methods). We used these pre-selected IRCC-PDX features as the input to 4 different lvl1 classifier pipelines: 1) model-based forward feature selection, followed by elastic net logistic regression, 2)

ANOVA-based feature selection, followed by either SVC or 3) extraTrees classifiers, and 4) a catBoost classifier pre-trained on a set of 55 multi-omic features from a collection of 860 pan-cancer cell lines from the Cell Model Passport portal (panCMP^45^), then refined on the same set of 55 features from the IRCC PDX (continual learning, Methods). The lvl1 predicted probabilities were then stacked and combined using a soft voting classifier which outputs a binary classification of cetuximab sensitivity (**Fig. 1bc**, Methods).

### Newly identified candidate biomarkers of cetuximab sensitivity

Our CeSta pipeline selects the most informative biomarkers of cetuximab sensitivity across training examples sampled from the IRCC-PDX collection by combining univariate statistical tests (Fisher’s exact, Mann-Whitney U test), percent lift, and logit (statsmodels v0.13.2 logit^46^) models (**Fig 1b, 1c, Tab S1**, and Methods). Here and in **Fig. 2a**, we provide an overview of some of CeSta’s top features (i.e. as ranked by their impact on CeSta’s predictions in **Fig 4a**) and their relationship with cetuximab sensitivity. The latter represents our binary target variable, with “responder” PDXs defined as those that grew in volume by 35% or less at three weeks after treatment (a proxy of disease control, as mentioned above) (**Fig 2b**, Methods).

Among our genomics features, beyond the KRAS-NRAS-BRAF triple negative signature, CLSPN (FET p = 0.05, percent lift: −0.675), PTEN (percent lift: −0.594), and PIK3CA (percent lift: −0.654) mutations were also more frequently observed in non-responder PDXs. Additionally, few other driver gene mutations such as EGFR (percent lift: −0.721) and MET (percent lift:- 0.702) were noticeably more likely to occur in non-responder PDXs, though rare overall (21 and 8 observations in IRCC-PDX, respectively). Only mutations in KRAS (logit p-value (logit p) = 0.002), BRAF (logit p = 0.037), PTEN (logit p = 0.049), and NRAS (logit p = 0.03) were found to be associated with cetuximab resistance via single-omic multivariate logit regression. Our CeSta approach combines these metrics (univariate and multivariate p-values, percent lift) into an aggregated feature selection score (Methods) which allows us to detect both well-supported and rare candidate markers.

CELLector subgroups 7 (APC, TP53, KRAS, PIK3CA mutated), 16 (TP53 wild-type; APC, KRAS, PIK3CA mutated), and 5 (APC, TP53, KRAS mutated; PIK3CA wild-type) were significantly associated with a non-responder phenotype (FET p = 0.002, 0.014, and 0.001, respectively). On the contrary, subgroup 12 (APC, TCF7L2, and TP53 mutated; KRAS, BRCA2, ATM, TPTE, EP400 wild-type) was approximately eight times (FET p = 0.011, percent lift: 7.934) more likely to contain responder PDXs. However, this was quite rare, with only 8 PDXs presenting this signature across IRCC-PDX. Subgroups 7,16, and 5 were also significantly associated with cetuximab resistance after multivariate logit regression (logit p = 2 x 10^−6^, 3 x 10^−6^, and 3 x 10^−4^, respectively).

Finally, FGFR1 CN gain events (FET p = 2 x 10^−4^, percent lift: 1.159) were more frequently observed in responder PDXs. Although ERBB2 and MET amplification events (i.e. more than two copies gained) were rare (5 and 3 examples in IRCC-PDX, respectively), they were more frequent in non-responders (percent lift: −1 for both).

These genomic signatures agree with previous surveys of CRC poor-prognosis driver alterations ^31, 47^, suggesting at least a partial overlap between markers of colorectal cancer progression and those of cetuximab resistance in PDX.

For transcriptomics features (**Fig 2c**), while EGFR (Mann-Whitney U test *p-value* (MWU p) = .4) and EGF (MWU p = .17) were not differentially expressed in cetuximab responders versus non-responders PDXs, REG4 (MWU p = 0.001), and EREG (MWU p = 7×10^−5^) were instead significantly upregulated in resistant and sensitive cases, respectively. REG4 (Regenerating Islet-Derived Protein 4) is a C-type lectin-like mitogenic protein known to stimulate EGFR signalling and promote migration and invasion in CRC^48^. High REG4 expression is associated with poor prognosis and low recurrence-free survival in CRC patients^49^ and, more specifically, with cetuximab resistance^50^ in CRC organoids and PDX models. A suggested mechanistic explanation points to FZD and LRP5/6, both upstream components of the Wnt/β-catenin pathway, which are involved in the REG4-mediated promotion of stemness induced by KRAS mutation in CRC with APC loss^51^. EREG (epiregulin) is a member of the EGF family and an EGFR ligand; it is thus involved in inflammation, cell proliferation, and cancer progression. EREG activity has been associated with cetuximab sensitivity in preclinical models and patients^52, 53^, and it has been suggested that, in an inflammatory environment, EREG can promote stemness and cancer cell proliferation by stimulating ERK signalling through EGFR activation in a variety of cancer types^54–56^.

We also observed high PROGENy^25^ EGFR pathway expression scores associated with a non-responder phenotype (MWU p = 0.002, percent lift: −1.879), whereas, as mentioned above, EGFR expression as an individual feature was not. We observed a similar pattern for KRAS: It was not differentially expressed across responders versus non-responders PDXs (MWU p = 0.23) but high MSigDB^57, 58^

HALLMARK_KRAS_SIGNALING_UP gene set ssGSEA scores were strongly associated with non-responder PDXs (MWU p = 0.001, percent lift: −10.688). These observations suggest that engineering aggregated expression features using ssGSEA and PROGENy scores might be more informative than individual gene expression features for cetuximab sensitivity prediction. However, it is also important to note that feature aggregation might introduce additional complexity. PROGENy signals for EGFR could be partly driven by downstream ERK-mediated signals, which are hard to disentangle from KRAS-triggered inputs. This may explain why both EGFRand KRAS signatures are associated with resistance to EGFR blockage.

Finally, we observed that higher MSigDB gene set ssGSEA scores for angiogenesis (percent lift: −2.168), inflammatory response (percent lift: −3.7), UV and DNA damage response (percent lift: −6.63), and Hedgehog (Hh) signalling (percent lift: −5.44), were all associated with non-responder PDXs (MWU p << 0.01 for all). The Hh hallmark score is fascinating as it might corroborate the evidence that Hh pathway activity correlates with reduced response to cetuximab^59^.

When considering methylation features (**Fig 2c**), NMF cluster 1, the most hypermethylated, was enriched for non-responders and MSI-like PDXs (FET p = 2 x 10^−4^, percent lift: −0.796). Cluster 4, the second-most hypo methylated, was strongly enriched for responder PDXs (FET p = 3 x 10^−4^, percent lift: 2.299).

Across all omics, both categorical (**Fig 2a**) and continuous features (**Fig 2b**) were either too sparse or too noisy to be adequate predictors of cetuximab response when considered individually. This highlights the effectiveness of an integrative model which combines the most informative features across ‘omic boundaries.

### Validation of the CeSta classifier

We set out to internally assess CeSta’s performance on our IRCC-PDX collection using a holdout shuffle approach, followed by testing the null hypothesis that results generated by different classifiers are equivalent^60^.

We started by generating 50 train/test set split (160 and 71 PDXs, respectively) replicates from our IRCC-PDX dataset. We used a nested cross-validation approach to tune and train 50 independent CeSta replicates (**Fig 1c,** “internal validation”). To provide a realistic and stringent benchmark, we evaluated many baseline cetuximab sensitivity classifiers of varying complexity (**Fig 3ab, S12**). Here, we present results from a performance comparison of our CeSta classifier against three of the best-performing baseline classifiers. These build on the SOTA clinical predictor of cetuximab sensitivity: the KRAS-NRAS-BRAF triple negative marker^39, 40^ and whether the original tumour is located in the left portion of the patient’s colon^37^. These features were combined into a cetuximab sensitivity classifier using either 1) a rule-based approach entirely analogous to the clinical criterion for cetuximab treatment (i.e. PDXs with the triple negative marker were predicted as responders to cetuximab, **Fig 3a**, “*tripleNegRule*” and “*tripleNegRightRule*”) or 2) an elastic net penalised logistic regression model (**Fig 3a**, “*elNet baseline*”) taking in input the four features above as possible regressors (Methods). As for CeSta, we tuned and trained 50 independent replicates of this latter baseline classifier over the 50 split replicates we previously generated.

**Figure 3.**
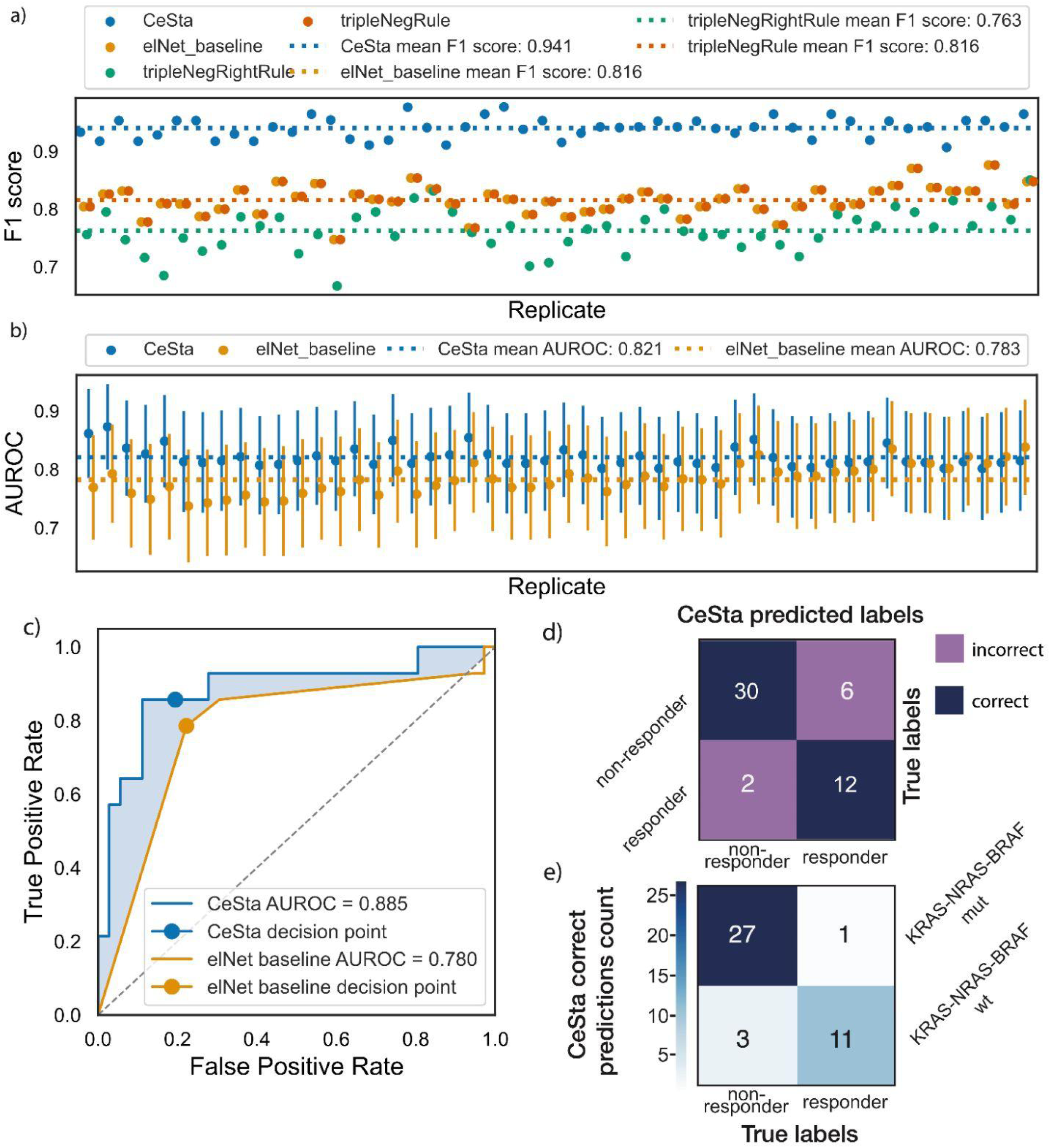
CeSta outperforms the state-of-the-art baseline classifier on IRCC-PDX and CR-PDX. **a)** Classification performances quantified through F1 scores (harmonic mean of precision and recall) across 50 train/test CR-PDX split replicates (x-axis) for the stacked classifier (“CeSta”, in blue), an elastic net penalised logistic model (‘elNet baseline’, in tan) which uses state-of-the-art clinical features for cetuximab sensitivity in CRC (KRAS, NRAS, BRAF mutational status, right colon tumour location), a rule-based classifier using the KRAS-BRAF-NRAS triple negative clinical signature (tripleNegRule, in orange) as a binary predictor, and another rule-based classifier which uses both the aforementioned triple-negative signature and the “right colon” feature (tripleNegRightRule, in green). **b)** Area under the receiver-operating-characteristic curve (AUROC) values and confidence intervals (vertical bars, obtained via DeLong’s method for the total error in the AUC^61, 62^) across 50 IRCC-PDX train/test split replicates (x-axis), for CeSta (in blue) and the elastic net penalised logistic model (‘elNet baseline’, in tan) described in a). **c)** AUROC (DeLong’s method) computed over the external validation CR-PDX dataset for CeSta (in blue) and the elNet baseline classifier (‘elNet baseline’, in tan) after both models are trained and tuned over IRCC-PDX. The orange-shaded area between the CeSta and elNet baseline ROC curves represents the improvement in AUROC. Decision point coordinates correspond to the false-positive and true positive rates obtained from the corresponding classifier’s predictions. Here, rule-based classifier decision points overlap with the elNet baseline’s. **d)** Confusion matrix from a comparison of CeSta classifier outcomes and PDXs actual cetuximab response over the external validation CR-PDX dataset. Correct predictions are on the diagonal highlighted in blue, incorrect predictions off the diagonal are highlighted in purple. **e)** CeSta correct prediction counts over the CR-PDX external validation set grouped by PDX cetuximab sensitivity (x-axis) and PDX KRAS-NRAS-BRAF triple-negative status (y-axis). CeSta correctly predicts additional triple-negative non-responders (3) and triple-positive responders (1), which all baseline classifiers miss.

**Figure 3** illustrates how CeSta outperforms all baseline models (mean F1: 0.941, Mann-Whitney post-hoc test pval: <<0.001) on this internal validation setup. Interestingly, the elNet baseline performance, measured via F1 score (i.e. the harmonic mean of precision and recall), fully matched the triple negative rule-based classifier, indicating that the elNet model can recapitulate the clinical decision criterion. **Figure 3b** shows that CeSta outperforms this same elNet baseline classifier for the vast majority of replicate splits (mean AUROC = 0.821 versus 0.780, Mann-Whitney post-hoc test pval: <<0.001), with an average .04 increase in ROC AUC, computed using the ROC AUC variance formula first proposed by Delong and colleagues^61–63^.

Following this encouraging result, we evaluated whether our CeSta classifier would outperform the clinical SOTA baseline classifier on an independent cohort of CRC PDX models (**Fig 1b,** “external validation”). This external validation cohort (from now on CR-PDX), consisting of 50 CRC xenografts, was collected and characterised at the genomic, transcriptomic and clinical levels at Charles River Discovery Research Services and included samples from European patients (Methods).

We tuned and trained CeSta and the baseline model over the entire IRCC-PDX collection (N = 231). We then compared their predictive performance on the never-before-seen CR-PDX set (N = 50) using the same set of multi-omic engineered features we described previously for IRCC-PDX (Methods). Similar to what we observed in the internal validation phase, our CeSta classifier outperformed the clinical baseline classifier (AUROC = 0.88 and 0.78, respectively), with an improvement of 0.1 ROC AUC (**Fig 3c**). More specifically, our CeSta pipeline correctly predicted three additional KRAS-NRAS-BRAF triple-negative PDXs as cetuximab non-responders and one additional triple-positive as a responder; on top of matching biomarkers correctly predicted by the baseline classifier (**Fig 3d,e**).

### Explanation of the CeSta classifier

Post hoc explanations approximate the behaviour of a classifier by modelling relationships between feature values and the classifier’s predictions. Here, we relied on SHapley Additive exPlanations (SHAP^64^) to define local feature importance and their impact on the CeSta classifier’s predictions. SHAP is a game theoretic approach through which values representing a feature’s average marginal contributions over all possible feature coalitions are computed.

Our CeSta classifier leverages additional informative genomic (e.g. FGFR1 amplification) and transcriptomics (e.g. EREG and REG4 expression; angiogenesis, inflammation, and Hh signalling ssGSEA scores) features (**Fig 4a**) to improve upon the clinical baseline classifier (**Fig 3b,c**) while retaining the latter’s top predictive features, namely the KRAS-NRAS-BRAF signature. As shown in the CeSta SHAP waterfall plot in **Figure 4b**, we observed high Hh signalling, high angiogenesis ssGSEA scores, and the KRAS, APC, TP53 mutation signatures being predictive of cetuximab resistance. In the same panel, high EREG expression and, more noisily, low REG4 expression and FGFR1 amplification appeared to influence the model towards a “responsive” prediction. Further, stacking our four lvl1 classifiers resulted in a slight performance increase over the best-performing lvl 1 classifier (i.e. the ANOVA SVC pipeline) taken on its own, albeit with substantial AUROC confidence interval overlap (**Fig 4e**).

**Figure 4.**
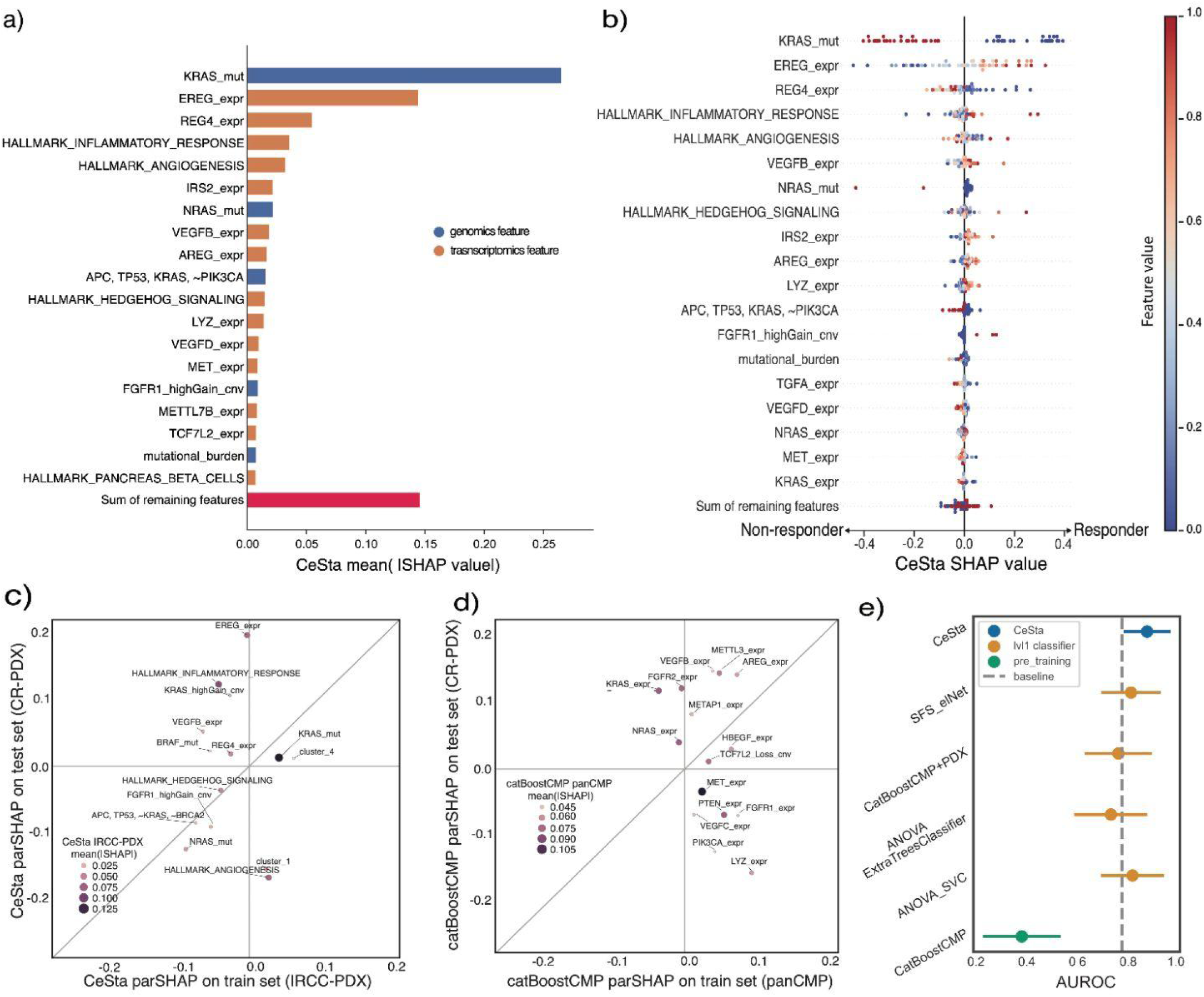
CeSta leverages informative features and combines weaker classifiers. **a)** Feature importance detected by CeSta, expressed as the mean absolute SHAP value (x-axis) for top significant features (y-axis). **b)** Feature impact on CeSta output using SHAP values (x-axis) for each top feature (y-axis) across all 50 PDXs in the CR-PDX validation set (scatter dots). The dot colour indicates the feature value in the corresponding example. The features with the largest importance in a) have the greatest impact on the model outcome, as well as a clean horizontal separation of positive and negative effects based on their value for that instance, as we can observe for KRAS mutation, EREG expression. **c)** CeSta top informative features’ performances on IRCC PDXs and the external cohort. Relationship between a CeSta feature’s SHAP values and cetuximab sensitivity on the train set (x-axis; full IRCC PDX set) and test set (CR PDX set) examples, after removing the effect of all other features (partial correlation, parSHAP). Dot size and colour indicate a feature’s mean absolute CeSta SHAP value on the training set, which quantifies a feature’s impact on model prediction (Fig 5a). The closer a dot is to the diagonal, the more its corresponding train and test parSHAP values are similar. CeSta’s top features (KRAS mutation, EREG expression, hedgehog signalling, NRAS mutation) all fall close to or above the diagonal, indicating either a very good fit on both datasets or slight underfitting. **d)** CMP-trained model’s top informative features underperform on the external cohort. Relationship between a CatBoostCMP feature’s SHAP values and cetuximab sensitivity on the train set (x-axis; panCMP set) and the test set (CR-PDX) after removing the effect of all other features (parSHAP). Dot size and colour indicate a feature’s mean absolute CatBoostCMP SHAP value on the training set, this quantifies a feature’s impact on model prediction. As for a), the closer a dot is to the diagonal, the closer its corresponding feature performs on our train and test sets. Many of this catBoost model’s top features (VEGFC expression, PTEN expression, MET expression) fall in the lower right quadrant of the plot, indicating overfitting. **e)** AUROC confidence intervals (CI, vertical bars, obtained via DeLong’s method) computed over the CR-PDX validation dataset for CeSta (in blue), three of the level 1 classifiers (in orange), a catBoost model trained on cell line data from the panCMP dataset (‘catBoost_panCMP’, in green), and the same catBoost model retrained on our IRCC-PDX dataset (‘catBoost_panCMP+IRCC-PDX’). Stacking four level 1 classifiers results in a slight performance improvement for CeSta over the best-performing level 1 classifier (ANOVA_SVC), albeit with substantial AUROC CI overlap. The cell-line trained CatBoost classifier is a poor predictor of cetuximab sensitivity in PDXs (no CI overlap). When this latter model is further trained on CR-PDX dataset (continual learning), its performance on the CR validation set is comparable with the other 3 IRCC-PDX trained lvl 1 classifiers.

We also detected very low collinearity among the top CeSta features’ values, with the strongest anticorrelation between the Hh signalling ssGSEA score and EREG expression (Pearson’s *r*: −0.2). In contrast, high EREG expression was associated with both increased angiogenesis (Pearson’s *r*: 0.4) and high inflammatory response (Pearson’s *r*: 0.3) ssGSEA scores (**Fig. S13**).

### Comparison of cetuximab response in cell lines and PDX models

PDX models are thought to recapitulate inter and intra-tumour heterogeneity observed in patients more faithfully than immortalised cell lines. They provide at least some stromal microenvironment interactions and are more likely to follow pathways of drug sensitivity or resistance found in primary human tumours^65^. However, 2d cell line models are undeniably cheaper as well as simpler to screen and characterise, an advantage that has enabled the generation of large multi-omics cell line datasets^45, 66, 67^ and aided systematic drug and functional genetic screening efforts^10, 66^.

We investigated whether a cetuximab sensitivity classifier trained 1) on a large pan-cancer multi-omic dataset (panCMP, N = 860) of 2d cell line models derived from the CMP dataset^45^, or 2) on a small colorectal cancer-specific subset of the same panCMP cell-line dataset (CRC-CMP, N = 44) would compare favourably against 1) the classifier itself, retrained on the IRCC-PDX dataset (N = 231) or 2) the classifier itself, retrained on a randomly selected subsample of IRCC-PDX, with the same size as the colorectal 2d cell-line dataset (subIRCC-PDX, N = 44).

We observed that a panCMP-trained boosting classifier (catBoost^68^ performed very poorly in predicting PDX sensitivity to cetuximab (**Fig. 4**). When this catBoost model was further trained on the IRCC-PDX dataset (continual learning, Methods), its performance on the CR-PDX validation set became comparable to that of the other IRCC-PDX trained lvl1 classifiers. We observed a similar result when we traded several examples for tissue specificity in the cell-line dataset and compared a CRC-CMP-trained classifier against itself after retraining on subIRCC-PDX (**Fig S14**).

We evaluated the partial correlation between a feature’s SHAP values and the target variable (parSHAP) to investigate further these differences in model performance across different training datasets. In this case, a positive parSHAP suggests that the classifier has identified and successfully exploited an informative feature for its current classification task. Given that our CeSta classifier performed just as well on the internal and external validations, it was not surprising to see matching parSHAP across CeSta SHAP values and cetuximab response in IRCC-PDX and CR-PDX (**Fig 4c**) for most features, and particularly for those with the most significant impact on model prediction (**Fig 4a,b**). On the other hand, several of the panCMP catBoost classifier’s top features (VEGFBC, PTEN, MET, PIK3CA and LYZ expression, TCF7L2 loss) did not perform as well on CR-PDX, compared to the cell lines training dataset (**Fig 4d**), that is: their SHAP values’ partial correlation with the target variable was lower across CR-PDX. This suggests that cell-line-trained models of cetuximab response struggle to predict PDX cetuximab sensitivity, primarily due to differences in the relationship between expression features and the target variable. These transcriptional differences between cell lines and PDXs might be due to the intense selection pressure imposed during cell line establishment, which makes available 2d models only partially representative of the general patient population^15^.

## Discussion

In this work, we describe and make available a multi-omic characterisation and drug screening data for one of the largest CRC PDX collections to date. This dataset recapitulates typical CRC alteration patterns observed in patient trials and gold-standard primary cohorts across all examined ‘omics, offering a precious combination of complete cetuximab response labels and dense multi-omic features. This cohort provides a realistic, stable platform for cetuximab sensitivity biomarker discovery and drug response modelling. Building on this PDX collection, we developed CeSta, a multi-omic ensemble classifier of cetuximab sensitivity based on a stacked ensemble architecture. When benchmarked against several state-of-the-art classifiers, CeSta outperformed all of them, including a classifier based on the current clinical criterion for patient stratification in cetuximab therapy prescription (KRAS, NRAS, BRAF negative mutational status and left sidedness of the primary tumour in the colon), both in a robust internal holdout shuffle validation and in a fully independent external validation dataset derived from an independent patient population.

CeSta identifies and leverages individual drug response biomarkers to increase cetuximab sensitivity prediction accuracy. These include epiregulin (EREG) and amphiregulin (REG4) expression levels, Hh signalling, angiogenesis, and inflammation gene set cumulative expression scores. These informative transcriptional features, in particular, show a weaker correlation with cetuximab response in 2d CRC models than PDXs, corroborating our observation of poorer predictive performance for models trained on cancer cell line datasets. The newly identified transcriptional biomarkers might be viable candidates for inclusion into an improved companion diagnostic for cetuximab sensitivity using clinical-grade gene expression technologies, such as Nanostring.

Collectively, our results illustrate the value of extensive, cancer type-specific, and well-characterised PDX collections for drug screening, drug sensitivity modelling and mechanism of action discovery, and motivate future efforts to increase resource dimensions and improve analytical approaches as a means to enhance further the informative power and translational potential of PDX-based research.

## Methods

### Primary samples

We set out to build a biobank of surgical materials from mCRC patients. The full study population consisted of tumour samples from 570 colorectal cancer patients that underwent surgical resection of liver metastases at the Candiolo Cancer Institute (Candiolo, Torino, Italy), the Mauriziano Umberto I Hospital (Torino), the San Giovanni Battista Hospital (Torino) and the Niguarda Hospital (Milano, Italy) from 2008–2015. Informed consent for research use was obtained from all patients at the enrolling institution before tissue banking, and study approval was obtained from the ethics committees of the centres. Tumour tissue (hepatic metastasis) not required for diagnosis was used to generate PDXs.

### Genomic data collection

Illumina PairEnd pre-capture libraries were synthesised from double stranded DNA according to Illumina’s protocol (Illumina Inc.). Genomic DNA was quality controlled and 200ng were used for library preparation per sample. DNA was sheared into 300 base-pair fragments (1ug DNA in 100ul volume) using the E210 Covaris plate system (Covaris, Inc. Woburn, MA). The fragmentation settings used are Intensity of 4, 200 Cycles per Burst, for 120 seconds. Sequencing libraries were amplified using the “bridge-amplification” process by Illumina HiSeq pair read cluster generation kits (TruSeq PE Cluster Kit v2.5, Illumina) and were hybridised to custom RNA baits for the Agilent SureSelect® protocol. Paired-end, 75bp sequence reads were generated using Illumina HiSeq 2000®. The sample mean sequencing coverage was ∼ 700X if the lost coverage because of duplicated and off-target reads is considered. The sequenced reads were aligned to the reference human genome (NCBI build37) using BWA-aln 0.5.9 (Li H. & Durbin R.,2009). The .bam files for all sequenced samples are stored at the European Genome-Phenome Archive (https://www.ebi.ac.uk/ega/ at the EBI) with accession number EGAD00001003334 (cram files are in EGAD00001003334, the study accession number is EGAS00001001171).

555 samples were sequenced using a custom-designed targeted colon cancer panel (SureSelect, Agilent, UK) consisting of all coding exons of 116 genes, 22 genes recurrently amplified/deleted, 51 copy number regions, 121 MSI regions and 2 gene fusions (RSPO2 and 3). Samples were fragmented to an average insert size of 150bp and subjected to Illumina DNA sequencing library preparation using Bravo automated liquid handling platform.

Sequencing was performed on an Illumina HiSeq2000 machine using the 75-bp paired-end protocol targeting 1Gb sequence per sample. Data quality was checked for 95% target coverage at 100x and mutation analysis was performed using an in-house algorithm. Sequencing reads are aligned to the NCBI 37 human genome build using the BWA algorithm^69^ with Smith-Waterman correction and PCR duplicates are removed. Base substitutions, small insertions or deletions, and breakpoints were identified by comparison against an unmatched control using established bioinformatic algorithms: CaVEMan (https://github.com/cancerit/CaVEMan/) for mutations, Pindel (https://github.com/genome/pindel) to detect insertions and deletions, and CNVKit (https://github.com/etal/cnvkit) for copy number detection.

We used an unmatched blood sample sequenced to an equivalent depth as control. To account for the absence of matched control, a bespoke variant selection pipeline was developed. To enrich for high-confidence somatic variants, we performed further filtering by removing: known somatic polymorphisms using human variation databases -- Ensembl GRCh37, 1000 genomes release 2.2.2 and ESP6500 -- and whether the same polymorphism was observed recurrently in 93 normal DNA samples sequenced using the same protocol and depth.

Cancer genes (CGs) are genes for which we can observe evidence of positive selection. Several statistical approaches have been developed to categorise the likelihood of a given gene in a specific tumour type to undergo a mutation at a high enough frequency for this to be indicative of a positive selection process. The majority of these methods rely on a comparison of non-synonymous (dN) and synonymous (dS) mutations in each gene and factor in additional covariates. We have elected to use as the foundation of our set of colorectal CGs two recent statistical approaches developed using large TCGA datasets ^70, 71^.

6426 driver variants across 113 genes are identified using the statistically significant single-codon hotspots from Chang *et a*l.^31^ and variants that are predicted as drivers using the intOGen^72^ framework. These variants are combined to generate a reference set of driver variants for this study, annotated based on their origin (Intogen driver only, Chang driver only, or common to both), their hotspot status, and whether they are known drivers of colorectal cancer. The final set of driver variants are used for annotating our PDX variants.

To assign segment log2R to individual genes we used coordinates overlap (BEDtools v2.29.2^73^, https://github.com/arq5x/bedtools2) between them and gene coordinates (TSS-TES) obtained from GENCODE (version 34, https://www.gencodegenes.org) for a set of 568 intOGen driver genes.

### TCGA COAD/READ copy number calling

We downloaded masked, segmented copy number variation (CNV) data from TCGA-COAD and TCGA-READ (∼1200 samples) on 02/09/2020 via the Genomic Data Commons Data Portal (GDC, https://portal.gdc.cancer.gov/repository) using the TCGAbiolinks R package (v2.20.0^74^).

The GDC CNV pipeline uses Affymetrix SNP 6.0 array data (harmonised to GRCh38) to identify genomic regions that are repeated and infer the copy number of these repeats. This pipeline uses the DNAcopy R-package^75^ to perform a circular binary segmentation (CBS) analysis. CBS translates noisy intensity measurements into chromosomal regions of equal copy number. The final output files are segmented into genomic regions with the estimated copy number for each region. The GDC further transforms these copy number values into segment mean values, which are equal to log2(copy-number/2). Diploid regions will have a segment mean of zero, amplified regions will have positive values, and deletions will have negative values ^76^. Masked copy number segments are generated using the same method except that a filtering step is performed that removes the Y chromosome and probe sets that were previously indicated to be associated with frequent germline copy-number variation.

We then ran both GISTIC2.0^77^ (ftp.broadinstitute.org/pub/GISTIC2.0] and ADMIRE v1.2^78^ (https://ccb.nki.nl/software/admire/) analyses across all the combined TCGA COAD/READ data to filter for “significant” copy number altered segments giving rise to robust CNV events across this patient cohort.

GISTIC2.0 was executed using the recommended “GISTIC2 Command Line Parameters” listed in the GDC copy number segmentation documentation at https://docs.gdc.cancer.gov/Data/Bioinformatics_Pipelines/CNV_Pipeline/#copy-number-segmentation. Here the “segmentation file” corresponds to the masked segmented copy number variation downloaded from TCGA COAD/READ, the “marker file” contains the aforementioned probe coordinates filtered for “freqcnv == FALSE” as per the GDC reference files (https://gdc.cancer.gov/about-data/gdc-data-processing/gdc-reference-files), and the “reference gene file” is the GRCh38 reference provided alongside GISTIC2.0 .

ADMIRE1.2 was executed using the same parameter configuration shown in the example use case provided at https://ccb.nki.nl/software/admire/readme.txt with the “segmented CNA” file again corresponding to the combined COAD/READ data, and the “marker file” containing the filtered probe coordinates.

The output of these two analyses identifies CNV events spanning multiple segments from different samples across the patient cohort. We then merged these results by computing the union of all (fully or partially) overlapping ADMIRE or GISTIC segments, and included all non-overlapping segments from either tool resulting in a set of 2382 events. From this combined output, we extracted event and segment coordinates and mapped both to 552 known cancer driver genes in the intOGen catalog^72^ (02/02/2020 release, https://www.intogen.org/download?file=IntOGen-Cohorts-20191112.zip) using BEDtools v2.29.2^73^ (https://github.com/arq5x/bedtools2).

Finally, we computed gene-specific CNV event frequencies by counting the number of TCGA samples with copy number altered segments mapping to both 1) the event and 2) the gene which shared the same CNV direction as the event, divided by the number of COAD/READ samples.

### Comparing driver gene SNPs in TCGA COAD/READ and PDXs

Frequencies of somatic alteration for TCGA samples was obtained from cBioPortal, selecting the Colorectal Adenocarcinoma TCGA, PanCancer atlas (https://www.cbioportal.org/study/summary?id=coadread_tcga_pan_can_atlas_2018) dataset.

### Comparing copy number variation events in TCGA COAD/READ and PDXs

We first binned PDX segment log2R values into three categories (“Loss”, “Neutral”, “Gain”), using the same GISTIC log2R thresholds we applied to the TCGA COAD/READ data (-.2, .1) [using the same threshold as in TCGA data here might be too strict for PDx sequencing data where there’s less non-tumour tissue contamination as murine cells/DNA are filtered out]. We then computed gene-specific CNV event frequencies by counting the number of PDX samples with copy number altered segments mapping to each gene, divided by the number of PDX samples.

We then computed the Spearman correlation coefficient for the TCGA and PDX genewise CNV event (here only “Loss”, “Gain”) frequencies.

### Assessing PDX copy number stability within lineages

We grouped 91 PDX samples, according to their genealogy, into 13 multi-passage lineages and retrieved gene-specific log2R data for 569 genes from the analysis described in the previous sections. We then computed the Pearson correlation across all gene log2Rs for each pair of PDX samples and labelled each Pearson coefficient according to whether the two samples belonged to the same lineage or to different ones.

### Assessing PDX mutational stability within lineages

We analysed somatic mutations along multi-passage PDX lineages using the same set of 91 PDX samples grouped into 13 lineages as described above. To rule out false positive calls for putative WT samples in lineages with apparent inconsistencies (Supplementary Figure S4), we further checked the coverage and absolute number of reads supporting each individual SNVs and only found single mutated reads in three WT samples with coverages ∼400X.

### Gene expression data collection

RNA was extracted using miRNeasy Mini Kit (Qiagen), according to the manufacturer’s protocol. The quantification and quality analysis of RNA was performed on a Bioanalyzer 2100 (Agilent), using RNA 6000 Nano Kit (Agilent). Total RNA was processed for RNA-seq analysis with the TruSeq RNA Library Prep Kit v2 (Illumina) following manufacturer’s instructions. Sequencing was then performed on Illumina Nextseq 500 at Biodiversa SRL, obtaining single end 151bp reads, aiming at 20M reads.

Read counts were obtained using an automated pipeline (https://github.com/molinerisLab/StromaDistiller), that uses a hybrid genome composed of both human and mouse sequences to exploit the aligner ability to distinguish between human derived reads, representing the tumour component, and mouse ones, representing the murine host contaminating RNA material.

Reads were aligned using STAR^79^ (version 2.7.1a, parameters --outSAMunmapped Within --outFilterMultimapNmax 10 --outFilterMultimapScoreRange 3 --outFilterMismatchNmax 999 --outFilterMismatchNoverLmax 0.04) versus this hybrid genome (GRCh38.p10 plus GRCm38.p5hg38 with GENCODE version 27 and mouse GRCm38 with GENCODE version 16, indexed with standard parameters and including annotation information from the GENCODE 27 plus m16 comprehensive annotation).

Aligned reads were sorted using sambamba^80^ (version 0.6.6) and only non-ribosomal reads were retained using split_bam.py^81^ (version 2.6.4) and rRNA coordinates obtained from the GENCODE annotation and repeatmasker track downloaded from UCSC genome browser hg38 and mm9. featureCounts (https://rdrr.io/bioc/Rsubread/man/featureCounts.html, version 1.6.3) was run with the appropriate strandness parameter (-s 2) to count the non multi-mapping reads falling on exons and reporting gene level information (-t exon -g gene_name) using combined GENCODE basic gene annotation (27 plus m16).

Sequencing data was available for 480 samples, but different filtering criteria lead to 470 QC passing samples. These criteria include: 1) >= than 15M total reads, 2) >= 60% reads assigned to genes by feature counts, 3) >= 30% reads assigned to human genes over the total of assigned reads.

These filters let us retain only samples with at least 5M human reads. Having defined the samples with acceptable mapping, we set out to identify the ones with lymphomatous characteristics^4^, to remove them. We considered different sources of information: 1) principal component analysis of the expression data itself (variance-stabilised); 2) computation of a sample level score for a leukocyte expression signature^82^, averaging the robust fpkm for all the signature genes; 3) methylation data, when available (see Methylation analysis); 4) histopathological analysis for a subset of tumours which were explanted and routinely stained with H&E.

This analysis highlighted a set of samples with lymphomatous characteristics pointed out coherently by the different data and prompted us to remove samples marked by at least one of the filters (specifically for expression PC2 >= 30 and leukocyte signature average >= 48), to correctly filter samples with only the expression data available.

Gene-level variance stabilised expression and robust fpkm values for 33670 genes were obtained using DESeq2^83^ (version 1.26.0), tmm using edgeR^84^ (version 3.28.1) using only read counts from human genes. CRIS and CMS subtyping was obtained for each individual tumour averaging the VST values for replicates, when available, using the R package CMScaller^34^ (v2.0.1, FDR = 0.05 and RNAseq = TRUE) and the R package CRISclassifier^4^ (v1.0.0, FDR< 0.2).

Github repositories: https://github.com/molinerisLab/StromaDistiller, https://github.com/vodkatad/RNASeq_biod_metadata and https://github.com/vodkatad/biodiversa_DE The .fastq files for all sequenced samples are stored at the European Genome-Phenome Archive (https://www.ebi.ac.uk/ega/ at the EBI) with accession number EGAS00001006492.

### Methylation data collection

Methylation profiles for 568 Colorectal Cancer samples were obtained using Illumina MethylationEPIC bead chip, which measures methylation status at about 850,000 sites using hybridization on two different probes after bisulfite treatment on DNA. These samples comprise tissue from the original patient, either primary tumours or metastases, or both in some cases, and the corresponding engrafted tumours in mice (PDXs). Raw data have been processed using the minfi package (https://bioconductor.org/packages/release/bioc/html/minfi.html, version 1.32.0). Data preprocessing was performed following the best practices outlined by Bioconductor minfi vignette and documentation, and Hinoue *et al*.^36^ (https://www.bioconductor.org/packages/devel/workflows/vignettes/methylationArrayAnalysis/inst/doc/meth <u>ylationArrayAnalysis.htm</u>l).

Background noise was removed using the minfi function *preprocessNoob()*, which implements the noob background subtraction method with dye-bias normalisation. Samples and probes that did not pass the quality control were then excluded from further analyses.

For samples, minfi provides a simple quality control plot that represents the log median intensity in both the methylated (M) and unmethylated (U) channels. By adopting the default median intensity cutoff of 10.5, six samples with lower values were removed from the dataset.

We then filtered the probes, based on their detection p-value (det-Pval), which is indicative of the quality of the signal. By filtering out all those probes of which det-Pval was higher than 0.01 in at least one sample, we removed 64,361 probes. We also removed all the probes mapping on X and Y chromosomes (19,627), to remove gender bias, and those probes that are known to bind to common SNPs (30,435). Moreover, using the list originally published by Chen *et al*.^85^, we removed 43,177 probes that have been demonstrated to map to multiple places in the genome.

To work with a coherent set of probes for all the samples, in particular xenografts, we decided to apply one last probes filter, removing all those probes known to specifically map on murine genome as well, in order to remove possible methylation signal coming from the murine infiltrate, with the same rationale followed for microarray data^82^. To do this, we combined two lists of murine-specific probes, obtained from Needhamsen *et al*.^86^ and Gujar *et al*.^87^, which resulted in removal of other 22,537 probes.

We combined the hg19 annotation package (IlluminaHumanMethylationEPICanno.ilm10b2.hg19 version 0.6.0), with the *liftOver()* function from the rtracklayer package^88^ (version 1.46.0) and the imported file hg19ToHg38.over.chain.gz (http://hgdownload.soe.ucsc.edu/goldenPath/hg19/liftOver/) in order to convert the remaining 700,298 probes’ coordinates from hg19 to hg38.

Moreover, as done for expression data (See Gene expression data collection), we removed samples with clear lymphomatous characteristics. Specifically for methylation, samples with PC2 >= 500 were almost always flagged by H&E analysis when it was available, therefore we considered all of them to be lymphomatous.

To identify groups of samples sharing similar methylation profiles, Beta values were used to run non-negative matrix factorization algorithms in R (https://www.rdocumentation.org/packages/NMF/, version 0.22.0). *k*=5 was identified as the best parameter by the cophenetic correlation coefficient (bootstrapping arguments: *rank=2:6, nrun=100, seed=42, .options=’p70’*). We therefore selected 5 as the number of classes used to characterise the methylation landscape of our samples.

The .idat files for all samples are stored at the Gene Expression Omnibus (GEO) with accession number GSE208713.

### Clinical data collection

Since the patients whose tumours are included in our biobank were not enrolled in a specific clinical trial and underwent surgery in different hospitals, our clinical data collection is based on personal communications with the Surgery Departments. This is the main reason behind the sparseness of the data.

### Measuring cetuximab response in PDX models

After surgical removal from patients, each metastatic colorectal cancer specimen was fragmented and either frozen or prepared for implantation: cut in small pieces and 2 fragments were implanted in 2 mice. After engraftment and tumour mass formation, the tumours were passaged and expanded for 2 generations until production of 2 cohorts, each consisting of 12 mice. Tumour size was evaluated once-weekly by calliper measurements and the approximate volume of the mass was calculated using the formula 4/3π·(d/2)^2^·D/2, where d is the minor tumour axis and D is the major tumour axis. PDXs derived from each original fragment were then randomised for treatment with placebo (6 mice) or cetuximab (6 mice) - animals with established tumours, defined as an average volume of 400 mm^3^, were treated with cetuximab (Merck, White House Station, NJ) 20 mg/kg/twice-weekly i.p.

For assessing PDX models response to therapy, we used averaged volume measurements at 3 weeks after treatment normalised to the tumorgraft volume at the time of cetuximab treatment initiation. 231 tumour grafts were classified as follows: 1) “objective response” (OR) models with a decrease of at least 50% in tumour volume 2) “progressive disease” (PD) models with at least a 35% increase in tumour volume, and 3) “stable disease” (SD) for the ones in between^2^.

Finally, to obtain a balanced dataset, we elected to combine the “SD” and “OR” classes into a single “SD-OR” (i.e. treatment responder) class, turning our cetuximab response modelling task into a binary classification problem.

All animal procedures were approved by the Ethical Commission of the Candiolo Cancer Institute and by the Italian Ministry of Health (authorization 806/2016-PR) All animal procedures for the CR PDX data set were executed in an AAALAC accredited animal facility and approved by the Committee on the Ethics of Animal Experiments of the regional council (Permit Numbers: G-13/13 & G18/12).

### Genomic feature engineering

To reduce data sparsity, we reshaped our mutational annotations into a binary matrix -- with columns (116 in total) corresponding to genes and rows (231 in total) corresponding to PDX models -- where a value of 1 indicates that one or more SNVs mapping to a given gene have been observed in a given PDX model. We also generated additional mutational features: a “mutational burden” feature containing the sum of all mutated genes for each PDX, and a set of “multiple mutations” features, indicating the number of unique SNPs hosted by a given gene in a PDX model. Finally, we filtered out any binary feature which was observed in fewer than 5 PDXs across our IRCC-PDX collection. To obtain a compact representation of relevant co-occurent or mutually-exclusive mutations, we developed an extended version of the CELLector methodology^27^ that partitioned the PDx mutation landscape recursively finding subgroups defined by the most recurrent combinations of genomic events (mutations or copy number alterations). Briefly, the original version of CELLector (from now on referred to as hierarchical), recursively applies the Eclat algorithm^89^ on a population described by a binary event matrix (BEM), with each column representing a genomic feature and 0/1 possible entries indicating the absence/presence of that feature in a sample. In the hierarchical version of CELLector the genomic background of a population is represented as a binary tree whose topology is defined by the most-frequently observed combination of genomic features (referred as signature) together with the fraction of samples for which those mutations occur and hence satisfy the signature rule (sequence of presence/absence of specific features). In particular, CELLector first identifies the root as the genomic feature with largest support, i.e. number of patients in which that feature is observed, and then defines two sibling nodes. The left child corresponds to the subset of samples satisfying the parent feature and the feature with greatest support among the samples in the parent node.

The right child corresponds to the complementary population of the parent node, composed of samples not satisfying that feature, and among those the feature with greatest support. This algorithm is applied recursively until no sub-population satisfying a certain signature rule of at least a *minGlobSupp* percentage of samples is identified, with *minGlobSupp* being a hyperparameter defined apriori. This hierarchical structure outputs K recursive signature rules that can be converted into a partition of K+1 groups as follows.

Starting from CELLector hierarchical binary tree,

1. for each node starting from the root, we define with U the set of samples satisfying that node rule defined as the corresponding signature S.
2. If the considered node has a left child (𝑈_𝑙_ ⊂ 𝑈) associated to feature 𝐹_𝑙_, we defined with 𝑈_𝑟𝑚_ : = 𝑈_𝑙_ the set of samples to be removed from U.
3. If 𝑈_𝑙_ has additionally a right child defined by feature 𝐹_𝑟_,𝑈_𝑟𝑚_ is updated with 𝑈_𝑟𝑚_: = 𝑈_𝑟𝑚_ ⋃𝑈_𝑟_
4. If 𝑈_𝑟_ has another right child 𝑈_𝑟,𝑟_ defined by signature 𝐹_𝑟,𝑟_,the update is repeated as 𝑈_𝑟𝑚_:= 𝑈𝑟𝑚 ⋃ 𝑈𝑟,𝑟 and this step is performed recursively until the considered node has no right child.
5. The new set of samples is defined as 𝑈_𝑛_ = 𝑈 \ 𝑈_𝑟𝑚_ and corresponding signature rule representing the group is defined as 𝑆, ∼ 𝐹_𝑙_, ∼𝐹_𝑟_, ∼𝐹_𝑟,𝑟_, …

If the condition in step 2. is not satisfied, the group is directly defined as samples in node U and satisfying signature S rule. Once every node in the hierarchical binary tree was considered, the last group is defined as the remaining samples that were not satisfying any hierarchical signature rule. The signature defining this group is created as the negation of the root node and all the recursive right childers, as described before. Note that the newly created groups could be composed of a fraction of patients lower than the *minGlobSupp*.

We applied the partitioned version of CELLector (V2.0.0) to the somatic mutation PDx space in BEM format with minGlobSupp fixed at 0.02.

Similarly to what we describe for above for mutation features, we discretise each of our 1163 gene-level log2 features into four categories (“Loss”, “Neutral”, “Gain”, “High Gain”), using, in addition to the GISTIC log2R thresholds for “Loss” and “Gain” (-.2, .1), an additional threshold at 2, above which a gene is considered to be involved in a “High Gain” event in which more than 1 additional copy is gained.

This “High Gain” category is added to help capture any association between driver gene high-order copy number gain and cetuximab sensitivity.

We then reshape these categorical copy number annotations into a binary matrix with columns corresponding to individual CNV events involving a given gene (e.g. “CD12_Gain”) and rows corresponding to PDX models. We then remove features which have the same value in 85% or more of our training PDX models.

After feature engineering and filtering, we are left with 1,295 genomic features.

### Transcriptomic feature engineering

To reduce RNAseq data dimensionality from an initial input of 33,670 gene-level expression features, as well as to include state-of-the-art knowledge of cancer signalling pathways and transcription factor activity, we computed 1) GSVA scores^90^ (http://www.biomedcentral.com/1471-2105/14/7) using the GSVA R package (version 1.34.0, R 3.6.3, kcdf=“gaussian”) on tmm expression levels and the MSigDB Hallmark gene sets^26^ as well as 2) PROGENy scores computed using the progeny R package^25^. Both sets of scores were computed separately for each train/test replicate (see following sections) to avoid any information leakage.

Finally, we considered that many PROGENy and Hallmarks gene set are partially overlapping: for example PROGENy’s “NFkB” set corresponds to Reactome’s “TAK1 activates NFkB by phosphorylation and activation of IKKs complex” and “RIP-mediated NFkB activation via ZBP1”, and thus it shares 8 of its 48 genes with PROGENy’s “TNFa” set (Reactome’s “TNF signaling”). To avoid excessive collinearity between scores based on overlapping gene sets, we first computed the Pearson correlation coefficient (PCC) for all pairs of engineered transcriptomic features over all instances in the training set, and considered as “collinear” all pairs with a PCC larger than .7. Here, for each pair of collinear features, we discard the one with the higher Mann-Whitney U test p-value between responder and non-responder PDXs. This yielded 38 engineered expression features.

### Clinical feature engineering

We consolidated our clinical data by: 1) dropping any features with more than 40% missing values, 2) dropping redundant or inconsistent features (“OXALIPLATIN-based treatments”, “N”, “T”,“N of other metastatic resections before collected metastasis”, “M”, “Site M”, “Site of primary”, “Site of primary DICOT”), 3) converting “Stage at first diagnosis” annotations to an integer score and retaining only the highest score for a given PDX model where multiple annotations are present, 4) converting the “ Lymph node density” annotations to a numerical score corresponding to the ratio of positive lymph nodes over the total lymph node count, 5) encoding all treatment backbone annotations as categorical features, 6) one-hot-encoding all sample anatomical location annotations. This yielded 25 features covering patient, previous treatment, and tumour metadata.

### Model architecture

For our cetuximab response model we selected a stacking classifier architecture. Stacking is an ensemble learning technique which combines the individual contributions of multiple classification models (level-1-classifiers) via a meta-classifier. Here, we use a soft voting classifier which outputs the final binary class labels (cetuximab non-responder; cetuximab responder) based on the argmax of the sums of the predicted probabilities from the level-1-classifiers (scikit-learn VotingClassfier^91, 92^, v1.02)

Our CeSta classifier uses a late integration approach to prevent high-dimensional ‘omics (transcriptomics, methylomics) from overwhelming smaller omics by dominating the selected feature set. We perform an initial round of single-omic supervised feature selection whose output is then piped into each of the four lvl 1 classifiers described below (**Fig. 1b**).

This selection step ranks features according to the product of 1) a feature rank based on the Fisher’s exact statistic (scipy v1.9^91, 92^) for binary features or Mann-Whitney U-test statistic (scipy v1.9) for continuous features, 2) a feature rank based on percent lift, and 3) a feature rank based on logit model (statsmodels v0.13.2 logit) coefficients. A set of top K features is then selected from this ranked list, with K being one of CeSta’s hyperparameters. This selection process is applied exclusively to the training set in each train, test split replicate during the internal validation (**Fig. 1c** and below) to avoid any information leakage. In Results and in Table S1 we are showing selected features and corresponding statistics and metrics obtained over the entire IRCC-PDX set as per the CeSta instance used for external validation (**Fig. 1c**)

We used 4 distinct level-1-classifier pipelines (**Fig 1b**): 1) a model-based (scikit-learn KNeighborsClassifier) forward feature selection, followed by elastic net penalised logistic regression (scikit-learn LogisticRegression with ‘penalty’ set to ‘elasticnet’), 2) ANOVA feature selection (scikit-learn f_classif), followed either by a support vector classifier (scikit-learn SVC) or 3) an extra trees classifier (scikit-learn ExtraTreesClassifier), and 4) a CatBoost classifier (catBoost 1.0.5^68^) trained on the same set of 30 features from CMP, then on IRCC PDX (continual learning).

Each level-1-classifier was trained (or re-trained in the case of CatBoost, see following sections) on a dataset of features selected (see above) from our 5 ‘omic data sources (mutation, CNV, expression, methylation, clinical). Finally, level-1-classifier prediction probabilities were stacked and taken as input by our meta-classifier (see above) which, in turn, gave in output a final binary prediction.

### Model training, tuning, and validation

We generated 50 train, test split (160/71 PDXs) holdout shuffle replicates by performing stratified sampling from our IRCC-PDX dataset. The latter consisted of 231 fully characterised (targeted sequencing, RNAseq, methylation assay, clinical metadata) PDX models which were labelled as cetuximab responders or non-responders according to tumour volume variation after treatment, as described above.

For the internal validation analysis, we used a nested cross-validation approach (inspired by mlextend’s StackingCVClassifier^93^) to tune and train 50 independent CeSta replicates, one per each train, test split. Each training set replicate was further split into 3 folds, and in 3 successive rounds, 2 folds were used (in turn) to fit the level-1-classifiers. In each round, the level-1-classifiers were then applied to the remaining 1 subset not used for model fitting in each iteration. The resulting predictions were then stacked and provided -- as input data -- to the meta-classifier. After comparing the meta-classifier’s prediction on the validation fold to the corresponding true labels, the first-level classifiers were fit to the entire training set replicate (**Fig 1c**).

This model training process was performed using a hyperparameter combination suggested by Optuna^94^ across 200 trials, while maximising the average of the area under the ROC curve (ROC AUC) computed over 3 training folds. Tuned parameter include: the number of top features selected during the first selection step, “colsample_bylevel”, “depth”, “boosting_type”,“boosting_type”,“bootstrap_type” for the CatBoost classifier; number of sequentially-selected features, elastic net ‘C’, ‘l1_ratio’ for the Logistic elastic net classifier pipeline; number of ANOVA-selected features, ‘C’ and ‘kernel’ for the SVC classifier pipeline; number of ANOVA-selected features, ‘n_estimators’ for the ExtraTrees classifier pipeline. This hyperparameter space search was performed, independently, for each model replicate.

Finally, we validated each of our 50 CeSta pipelines by predicting each PDX model in their respective test set as a cetuximab “responder” or “non-responder”, and computing the resulting ROC AUC and ROC AUC .95 confidence interval (using DeLong’s method) by comparing predicted and true labels.

For the external validation analysis, the same tuning, training, and validation process was repeated using the entire IRCC-PDX dataset as a training set (N=231), and the CR-PDX dataset as a test set (N=50).

### Performance baselines

To provide a realistic benchmark for CeSta performance, we define and train a number of alternative, multi-omic cetuximab sensitivity predictors. The latter are all trained, tuned, and validated using a set of 30 holdout shuffle replicates, analogous to the setup we use for CeSta internal validation in Figure 1c.

“*tripleNegRule*” is a rule-based classifier based on the KRAS-NRAS-BRAF mutational signature: it will output a “non-responder” prediction if any of these three genes is mutated in the current PDX example.

*tripleNegRightRule”* is a rule-based classifier based on the KRAS-NRAS-BRAF mutational signature and the “right colon” marker (i.e. whether the original tumour was located in the right portion of the patient’s colon). This decision strategy originates from a retrospective analysis of triple negative patients from the CRYSTAL and FIRE-3 trials where right-sided tumours had significantly poorer prognosis and lower response to cetuximab treatment^37^.

*tripleNegRightRule* will output a “non-responder” prediction if either 1) any of KRAS, NRAS, BRAF is mutated or 2) the original tumour was right-sided.

“*elNet_baseline*” is an Elastic-Net net penalised logistic regression classifier (scikit-learn LogisticRegression with penalty set to “elasticnet”) based on 4 binary features encoding the mutational status of KRAS, BRAF, NRAS (i.e. the ‘triple negative’ CRC signature), and whether the primary tumour is located in the Right Colon. This corresponds to the state-of-the-art clinical signature for cetuximab sensitivity in colorectal cancer, as we discuss in Introduction and Results.

“*rawL1elasticnet*” is an Elastic-Net net penalised logistic regression classifier which uses our full set of 39960 raw (non-engineered, non pre-selected) features, that is: binary gene mutational status features, variance-normalised gene-level RNAseq data, clustered methylation probe signal, and binary CNV events.

*MixOmics sPLS-DA”* uses mixOmic’s^95^ multivariate integration approach, based on Partial Least Squares (PLS) regression and discriminant analysis, in which the most informative features (i.e. those that best discriminate between cetuximab responsive and non-responsive PDXs) from different ‘omics are selected with the constraint of correlation between their first PLS components. More specifically, here we follow the multi-omic classification case study illustrated in http://mixomics.org/methods/spls/. We 1) perform LASSO feature selection (glmnet v4.2, https://www.rdocumentation.org/packages/glmnet) for methylation (700,298 non-engineered features) and expression (33,670 non-engineered features), 2) use a sparse partial least-squares discriminant analysis model (sPLS DA) for single-omic dimensionality reduction, 3) followed by a DIABLO model for horizontal multiple ‘omics integration. We optimise both the number of PLS components and the number of selected features for each omic and each component via 3-fold cross validation on each training set replicate.

Finally, we validate these benchmark classifiers on each test set replicate, as described for our CeSta classifier in **Fig 3b** by labelling each PDX model as a cetuximab “responder” or “non-responder”, and computing the resulting ROC AUC by comparing predicted and true labels, again using DeLong’s method for computing the ROC AUC 0.95 confidence interval where possible.

### Cell Model Passport datasets

The Cell Model Passport portal^45^ (https://cellmodelpassports.sanger.ac.uk/) catalogues and curates multi-omic data for cancer cell line and organoid models. When combined with the Genomics of Drug Sensitivity in Cancer (GDSC) dataset (https://www.sanger.ac.uk/tool/gdsc-genomics-drug-sensitivity-cancer/), it provides genomics, transcriptomics, and cetuximab response data for 860 unique cancer cell line models (panCMP dataset). Here, we repeat the same data preprocessing and feature engineering steps we performed for the IRCC-PDX dataset, with the exception of the NMF-based clustering of methylation probes as this omic is missing from the CMP collection. Further, as cell line cetuximab response is quantified as IC50 values, rather than tumour volume change, here we dichotomise our target variable using the median IC50 for all cell lines in the panCMP dataset with lines falling below this threshold being labelled as “responders”.

For the purpose of comparing the predictive performance of a model trained on cell line data against one trained on PDX data, we generate a panCMP training set which includes a subset of 860 examples and 55 expert-selected multi-omic features (Data and Code Availability). This feature subset is available in both the aforementioned panCMP dataset, our IRCC-PDX dataset, and the CR-PDX dataset. This feature subset fully overlaps with available features from our IRCC PDX train set. We then train and tune a catBoost classifier pipeline (see above for pipeline architecture, hyperparameters space) over this panCMP training set using an 8-fold cross-validation approach across 50 Optuna trials. This cell-line trained “base model” is then provided, as a starting point for continual learning, to a second round of training (using the ‘init_model’ parameter) over either an IRCC PDX train set replicate for internal validation, or the entire IRCC-PDX dataset for external validation on the CR-PDX dataset.

From the panCMP dataset, we can further subset 44 colorectal cell lines (CRC-CMP), which are characterised with the same set of 55 features as in the panCMP dataset. This context-specific dataset can also be used to train a catBoost “base model” which we then feed into a second catBoost classifier trained on IRCC-PDX.

### External validation: Charles River dataset

An independent CRC PDX cohort^96^ (https://www.cancermodels.org/data/?facets=model.data_source:CRL%20AND%20model.model_type:xenograft%20AND%20patient_tumour.cancer_system:Digestive%20System%20Cancer) has been assembled and characterised by our collaborators at Charles River Discovery Research Services (CR). We use 50 CRC LMX, first-pass PDX models corresponding to 50 unique patient samples characterised using the same set of multi-omics as in the IRCC PDX cohort. For missing features (e.g. methylation cluster labels) we impute their values for this cohort using the mode for categorical features and the median for continuous features. We then use this CR-PDX dataset as a fully independent validation cohort to compare our stacked classifier’s performance against that of baseline models after training on the entire IRCC-PDX dataset.

### Post-hoc model explanation

As a cross-model proxy for feature importance, for each feature, we calculate the mean of the absolute SHAP values (https://github.com/slundberg/shap v0.4) across all instances in the test set. We consider the absolute values as we do not want positive and negative values to offset each other. Features that have large mean absolute SHAP values are those that more significantly impact model predictions.

We are also interested in assessing, for a given classifier, which features perform equally well across different datasets (i.e. panCMP, IRCC-PDX, CR-PDX). To do so, we start by evaluating the relationship between a feature’s SHAP values and the target variable. A positive correlation here indicates that the model has identified and it is successfully exploiting an informative feature for its current classification task. Given that SHAP values are additive, with the model’s prediction being the sum of all feature SHAPs, it makes sense to remove the effect of other features’ contribution by computing the partial correlation between each feature and the target after removing the effect of all other features (i.e. controlling variables). Specifically, here we use pingouin^97^ (v0.5.1) and its partial_corr function specifying, in turn, all features but one as x-covariates.

### Data and code availability

https://github.com/molinerisLab/StromaDistiller and https://github.com/vodkatad/RNASeq_biod_metadata contain a pipeline tracking counts and metadata across different sequencing batches for xenografts/organoids RNAseq.

https://github.com/vodkatad/biodiversa_DE contains code to perform DEG analyses with DESeq2 and various enrichment analyses on results.

Code for multiomic data preprocessing and integration is available at https://bitbucket.org/uperron/pdx_multiomics_integration_preproc

Code and data to fully replicate the results in Figures 1,2 is available at https://bitbucket.org/uperron/ircc-pdx_exploration

Code, models, and data to fully replicate CeSta and the results in Figures 3,4 is available at https://bitbucket.org/uperron/cesta_pdx

CELLector is available at https://github.com/francescojm/CELLector.

*Raw sequencing data:*

1. Targeted DNA sequencing: https://ega-archive.org/studies/EGAS00001001171
2. RNAseq: https://ega-archive.org/studies/EGAS00001006492

Access to these datasets will be granted upon request via the EGA portal, as required for personally identifiable data.

*Methylation data:*

1. https://www.ncbi.nlm.nih.gov/geo/query/acc.cgi?acc=GSE208713

We set up a temporary token to access this dataset during review: cfihygswjlohxqd. This data will be made fully public upon publication of this manuscript.

### Author contribution

U.P., E.G., A.C., N.C., U.M., L.Tru., A.B., and F.I. conceived the project and scope.

E.R.Z. derived and characterised the IRCC-PDX collection, collected metadata from the original patients, coordinated by E.M., M.E., L.Tru., and A.B.

U.P., E.G., A.C., M.V., E.K., L.Tra., C.I., and I.M. processed and analysed the IRCC-PDX data. U.M., L.Tru., A.B., and F.I. supervised IRCC-PDX data processing and analysis.

H.K. and J.S. derived, characterised, and analysed the CR-PDX collection.

U.P. designed and implemented the CeSta pipeline. U.P., E.G., and A.C. drafted the manuscript and designed the figures. U.P., E.G., A.C., L.Tru., A.B. and F.I. edited and revised manuscript and figures. All authors discussed the results and contributed to the final manuscript.

