## Supplementary Information for "Integrative ensemble modelling of cetuximab sensitivity in colorectal cancer PDXs"

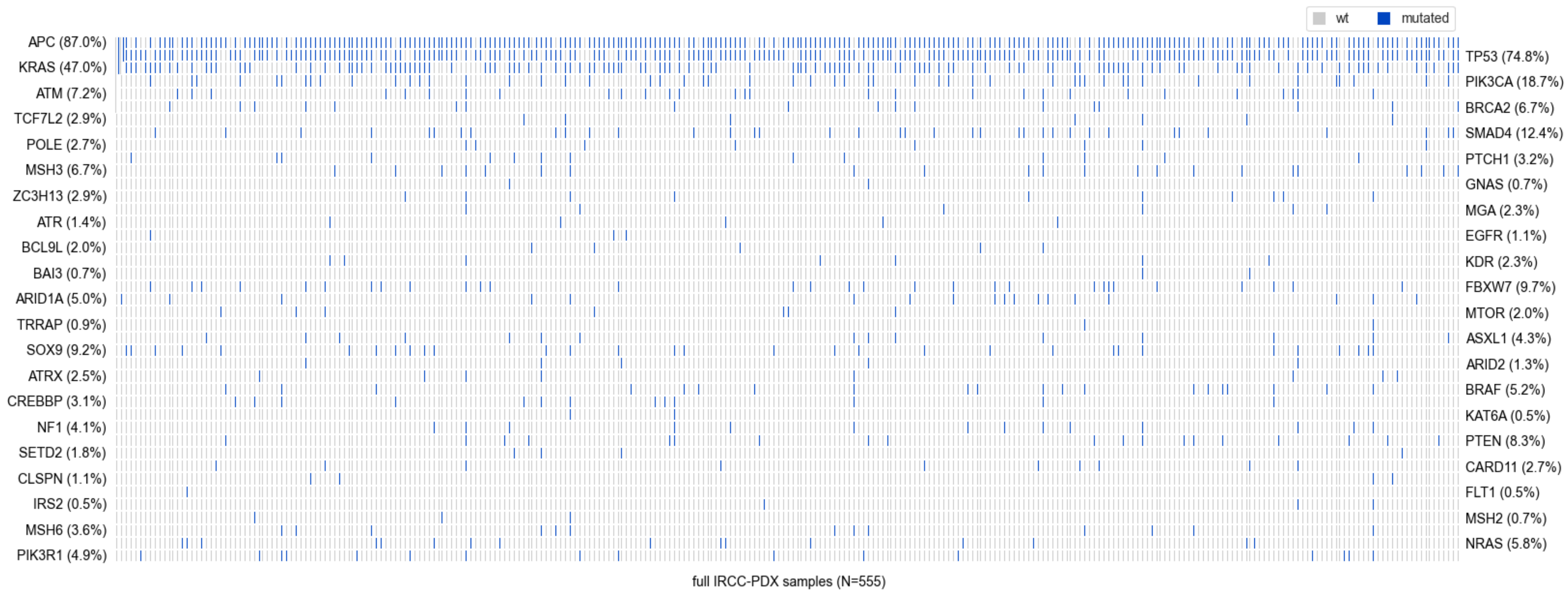
wt
mutated
APC (87.0%)
KRAS (47.0%)
ATM (7.2%)
TCF7L2 (2.9%)
POLE (2.7%)
MSH3 (6.7%)
ZC3H13 (2.9%)
ATR (1.4%)
BCL9L (2.0%)
BAI3 (0.7%)
ARID1A (5.0%)
TRRAP (0.9%)
SOX9 (9.2%)
ATRX (2.5%)
CREBBP (3.1%)
NF1 (4.1%)
SETD2 (1.8%)
CLSPN (1.1%)
IRS2 (0.5%)
MSH6 (3.6%)
PIK3R1 (4.9%)
TP53 (74.8%)
PIK3CA (18.7%)
BRCA2 (6.7%)
SMAD4 (12.4%)
PTCH1 (3.2%)
GNAS (0.7%)
MGA (2.3%)
EGFR (1.1%)
KDR (2.3%)
FBXW7 (9.7%)
MTOR (2.0%)
ASXL1 (4.3%)
ARID2 (1.3%)
BRAF (5.2%)
KAT6A (0.5%)
PTEN (8.3%)
CARD11 (2.7%)
FLT1 (0.5%)
MSH2 (0.7%)
NRAS (5.8%)
full IRCC-PDX samples (N=555)

***Figure S1 - Top frequently mutated genes in the full IRCC PDX cohort (see Methods)****. For each of 555 IRCC-PDX models (x-axis) the heatmap indicates whether one or more mutations are observed in a given gene (y-axis).*

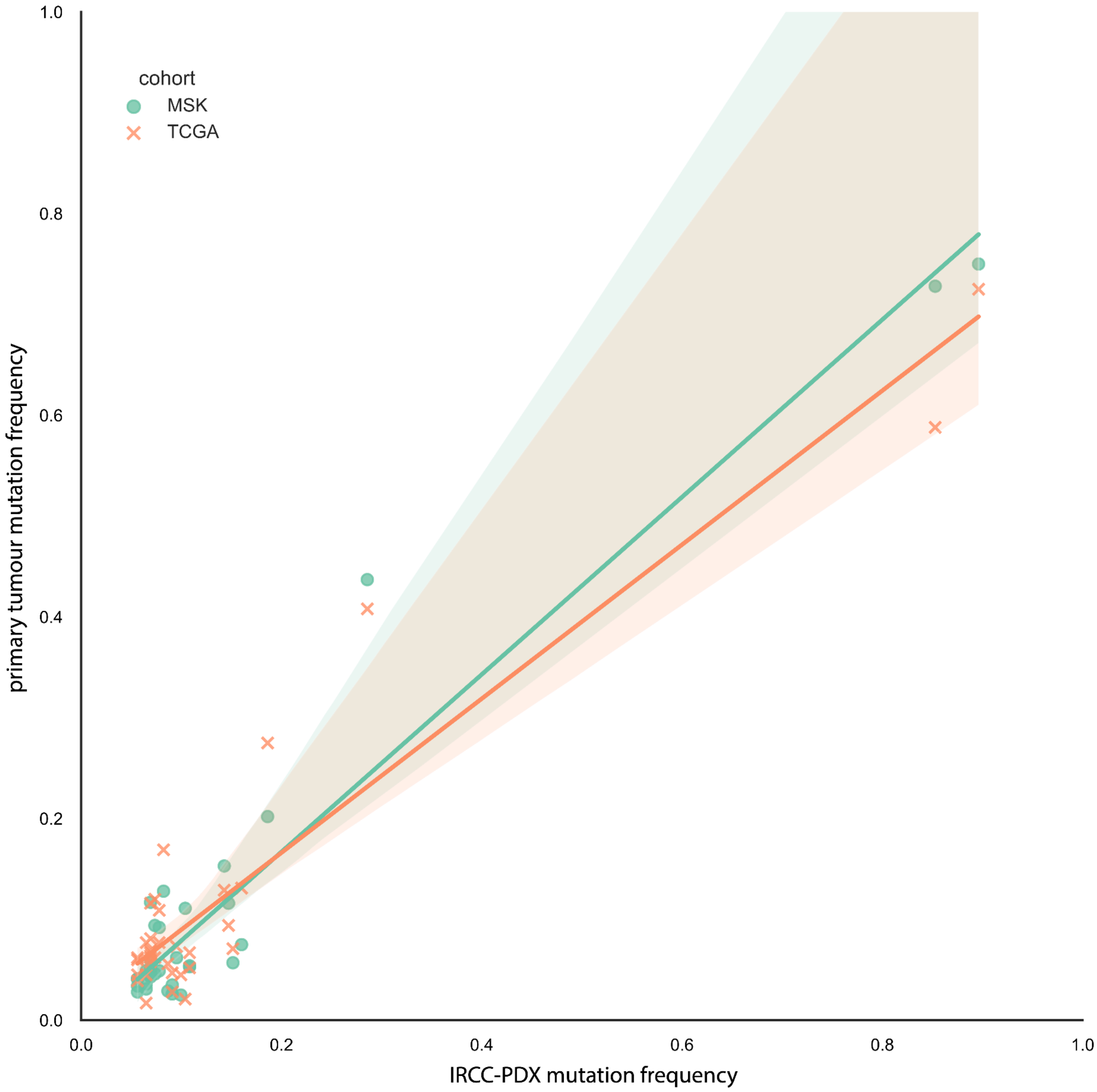


***Figure S2 - Mutation frequencies observed in TCGA COAD/READ and MSK-IMPACT, are conserved in IRCC-PDXs.***

*Strong positive correlation between gene mutation frequency in TCGA COAD/READ (Spearman r: 0.51,in orange), MSK-IMPACT (Spearman r: 0.625, in green) on the y-axis, and IRCC-PDX (on the x-axis) for the top 70 most frequently mutated genes.*

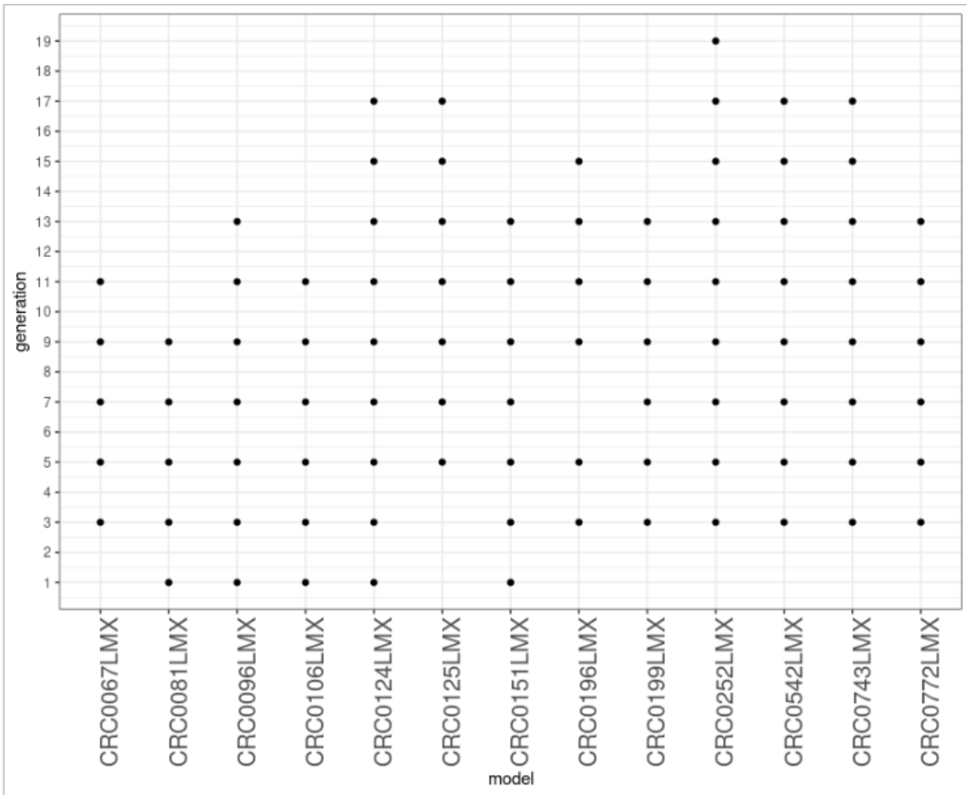


***Figure S3 - multi-passage lineages in IRCC-PDX.*** *Summary of the genealogy of 91 PDX samples grouped by generation (on the y-axis) and lineage (on the x-axis). For most models, odd passages between 3 and 13, are available.*

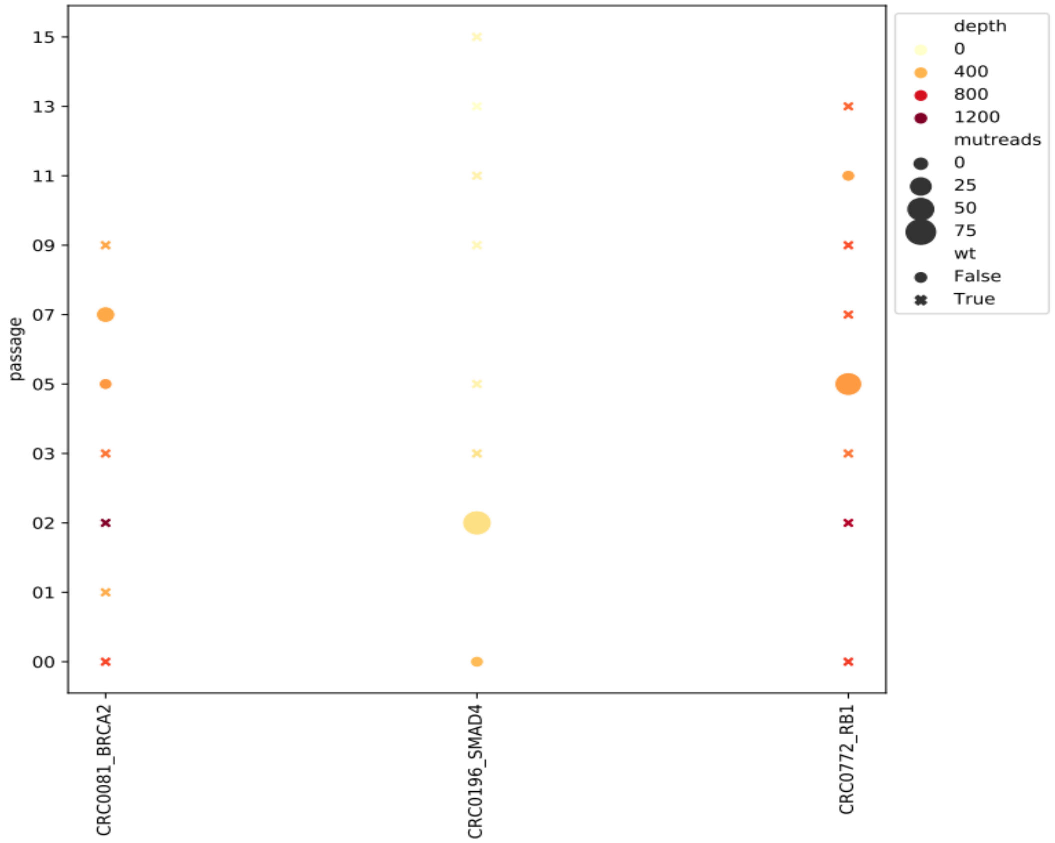


***Figure S4 - SNP anomalies within PDX lineages.*** *Mutational status (point shapes), sequencing read count (colour shades), and percentage of mutated reads (point sizes) for each available PDX sample (on the y-axis) along 3 multiple-passage lineages (on the x-axis) containing SNP anomalies. These mutational patterns support the existence of rare detectable subclones harbouring mutations in single samples across xenopatients lineages.*

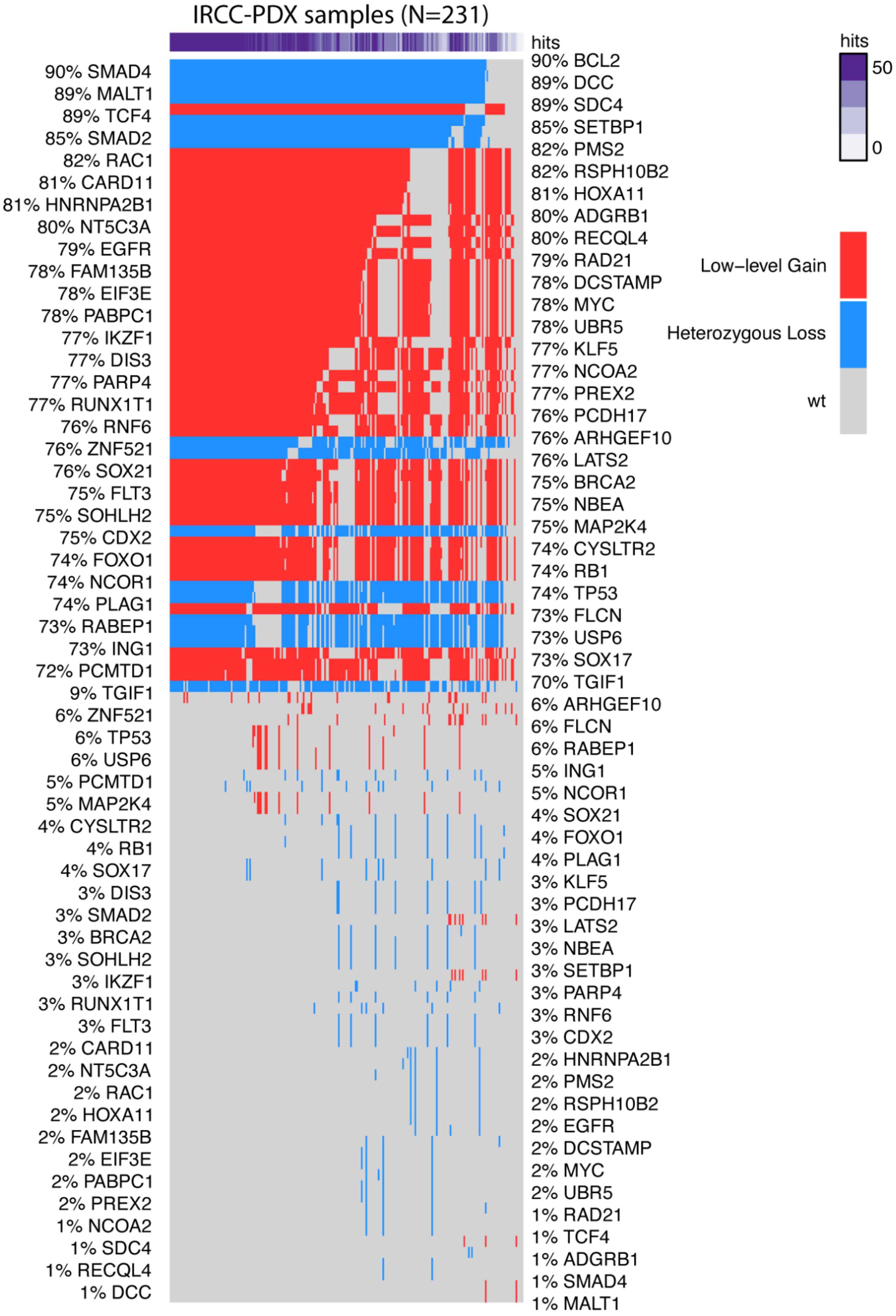
IRCC-PDX samples (N=231)
hits
90% SMAD4
89% MALT1
89% TCF4
85% SMAD2
82% RAC1
81% CARD11
81% HNRNPA2B1
80% NT5C3A
79% EGFR
78% FAM135B
78% EIF3E
78% PABPC1
77% IKZF1
77% DIS3
77% PARP4
77% RUNX1T1
76% RNF6
76% ZNF521
76% SOX21
75% FLT3
75% SOHLH2
75% CDX2
74% FOXO1
74% NCOR1
74% PLAG1
73% RABEP1
73% ING1
72% PCMTD1
9% TGIF1
6% ZNF521
6% TP53
6% USP6
5% PCMTD1
5% MAP2K4
4% CYSLTR2
4% RB1
4% SOX17
3% DIS3
3% SMAD2
3% BRCA2
3% SOHLH2
3% IKZF1
3% RUNX1T1
3% FLT3
2% CARD11
2% NT5C3A
2% RAC1
2% HOXA11
2% FAM135B
2% EIF3E
2% PABPC1
2% PREX2
1% NCOA2
1% SDC4
1% RECQL4
1% DCC
90% BCL2
89% DCC
89% SDC4
85% SETBP1
82% PMS2
82% RSPH10B2
81% HOXA11
80% ADGRB1
80% RECQL4
79% RAD21
78% DCSTAMP
78% MYC
78% UBR5
77% KLF5
77% NCOA2
77% PREX2
76% PCDH17
76% ARHGEF10
76% LATS2
75% BRCA2
75% NBEA
75% MAP2K4
74% CYSLTR2
74% RB1
74% TP53
73% FLCN
73% USP6
73% SOX17
70% TGIF1
6% ARHGEF10
6% FLCN
6% RABEP1
5% ING1
5% NCOR1
4% SOX21
4% FOXO1
4% PLAG1
3% KLF5
3% PCDH17
3% LATS2
3% NBEA
3% SETBP1
3% PARP4
3% RNF6
3% CDX2
2% HNRNPA2B1
2% PMS2
2% RSPH10B2
2% EGFR
2% DCSTAMP
2% MYC
2% UBR5
1% RAD21
1% TCF4
1% ADGRB1
1% SMAD4
1% MALT1
hits
50
0
Low−level Gain
Heterozygous Loss
wt

***Figure S5* - *Copy number alteration landscape in IRCC-PDX.*** *Most frequently copy number (CN) altered genes (y-axis) across all 231 PDXs in the IRCC-PDX cohort (x-axis). In the rug plot at the top, colour scheme indicates the total CN alteration ("hits") count for a given PDX model (darker stands for higher counts). In the main heatmap, colours separate CN alteration events by direction: loss (in blue) and gain (in red), with grey indicating no CN alterations. Y-axis annotations report CN alteration frequency for a given gene across our IRCC-PDX dataset.*

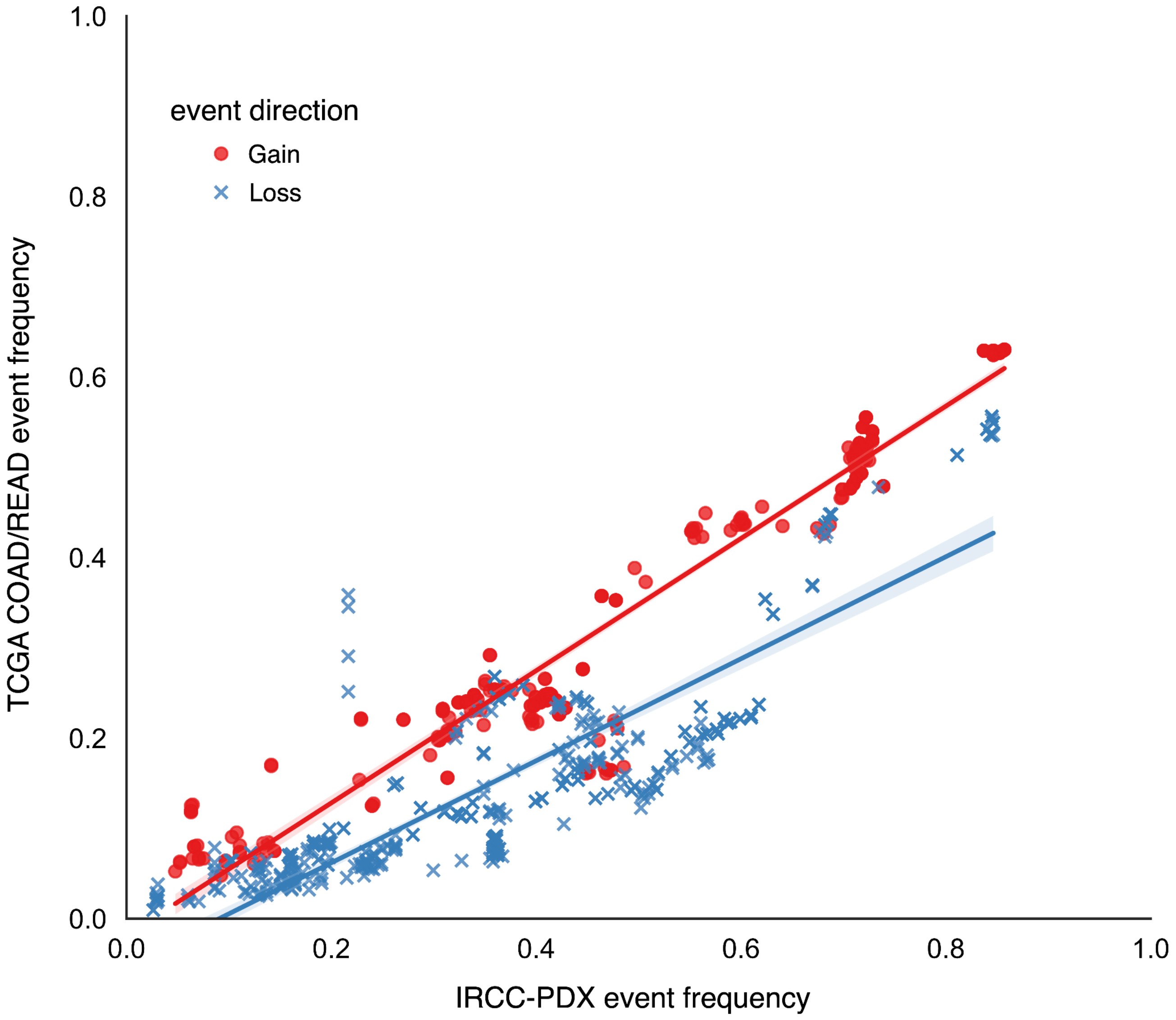


***Figure S6* - *Gene copy number variation event frequency observed in TCGA COAD/READ samples is conserved in IRCC-PDX samples.*** *Strong positive correlation (Spearman r Loss: 0.934; Spearman r Gain: 0.875) between the TCGA COAD/READ (on the y-axis) and IRCC-PDX (on the x-axis) gene copy number*

*variation event frequencies across 548 genes, grouped by event direction in TCGA (Loss in blue, Gain in red).*

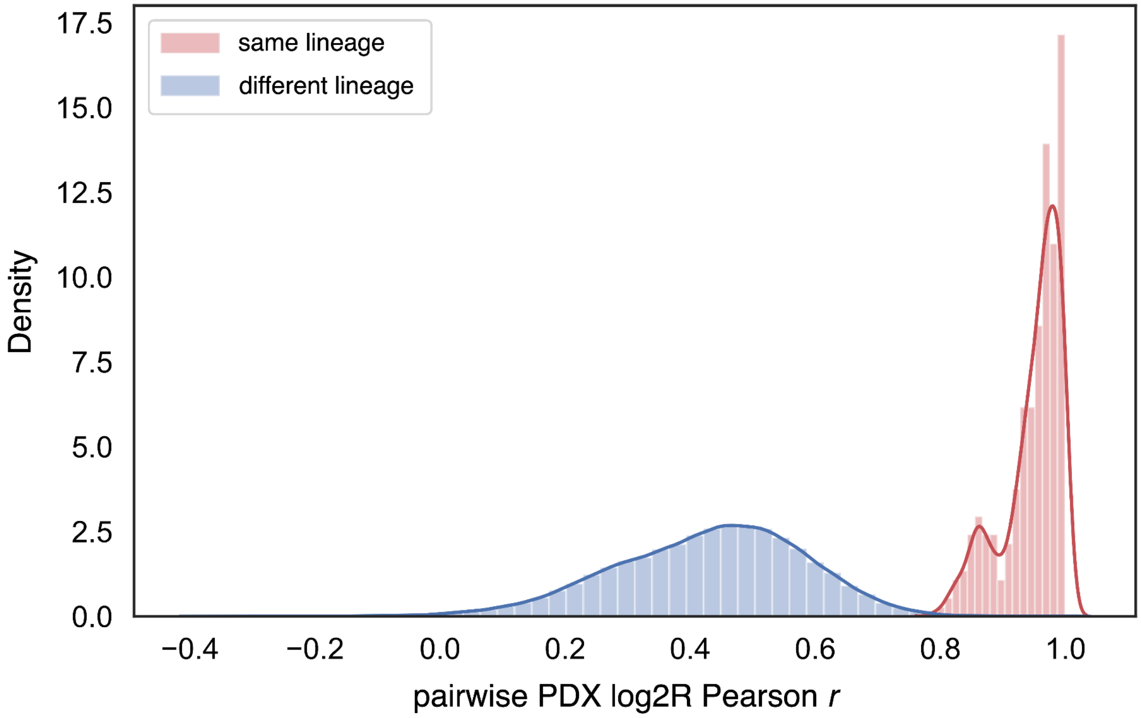


***Figure S7 - Gene log2R is strongly correlated among PDX samples belonging to the same lineage.*** *Distributions of Pearson's correlation coefficients (Pearson r, on the x-axis) computed across 557 gene $log_2$ ratio copy number values (log2R) for all pairs of 91 PDX samples belonging to one of 13 multiple-passage lineages, grouped according to their PDX lineage (as indicated by the different colours). Intra lineage comparisons yield much higher Pearson r values (median: 0.927) than inter lineage comparisons (median: 0.376).*

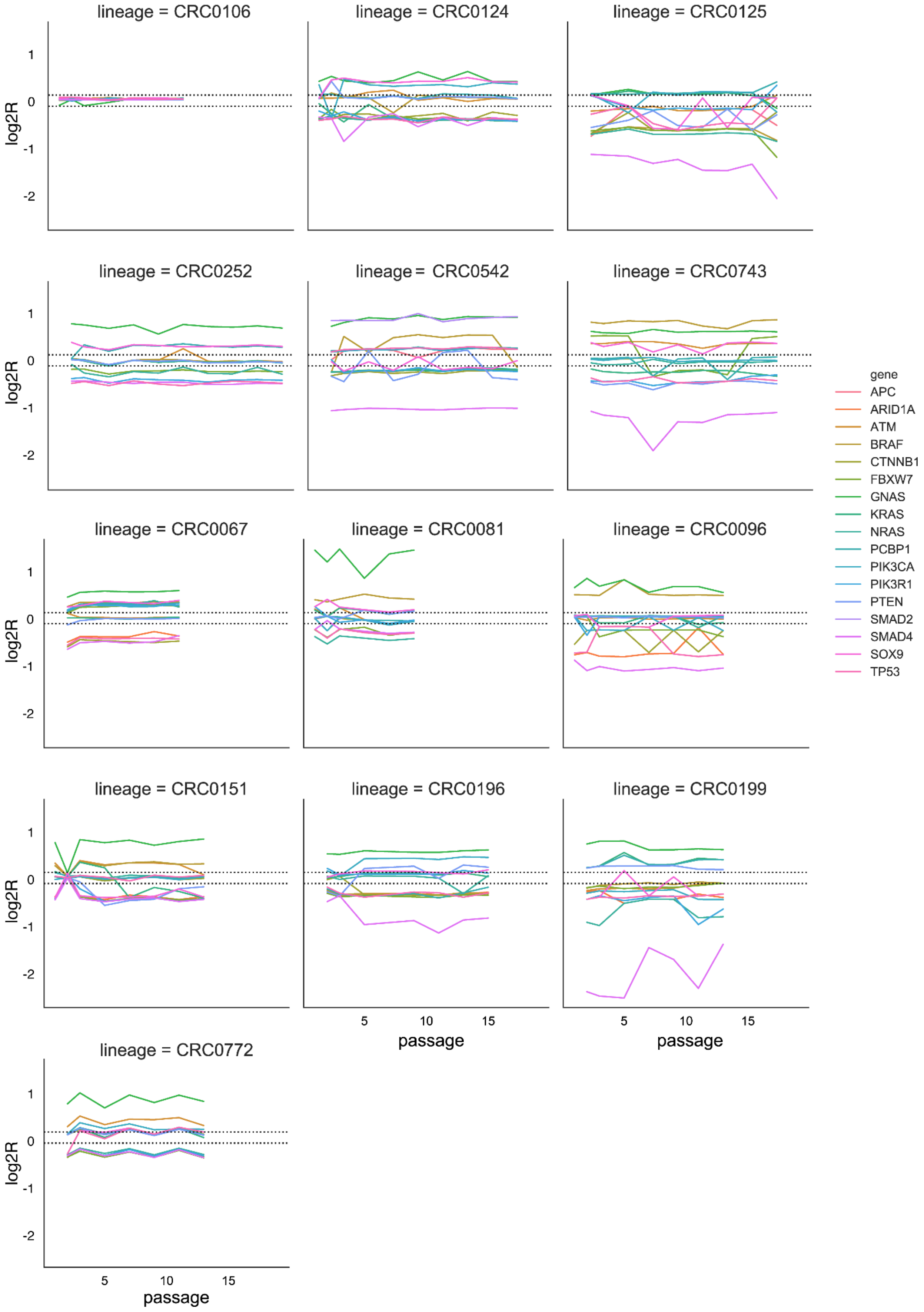

lineage = CRC0106
lineage = CRC0124
lineage = CRC0125
lineage = CRC0252
lineage = CRC0542
lineage = CRC0743
lineage = CRC0067
lineage = CRC0081
lineage = CRC0096
lineage = CRC0151
lineage = CRC0196
lineage = CRC0199
lineage = CRC0772
log2R
1
0
-1
-2
5
10
15
passage
gene
APC
ARID1A
ATM
BRAF
CTNNB1
FBXW7
GNAS
KRAS
NRAS
PCBP1
PIK3CA
PIK3R1
PTEN
SMAD2
SMAD4
SOX9
TP53

***Figure S8 - Driver Gene copy numbers are stable within PDX lineages.*** *Gene-specific log2R copy number values (on the y-axis) across passages (on the x-axis) across 13 multiple-passage PDX lineages (one per plot), considering 17 cancer driver, CRC-associated genes (Methods) hosting copy number alterations in at least one lineage. Each curve corresponds to a gene, the two horizontal dotted lines indicate "Gain" (.1) and "Loss" (-.2) GISTIC log2R thresholds, respectively. Across these lineages,the majority of genes (median: 94.112 %) remains within the same y-axis bin (i.e. gene copy number is constant) across passages.*

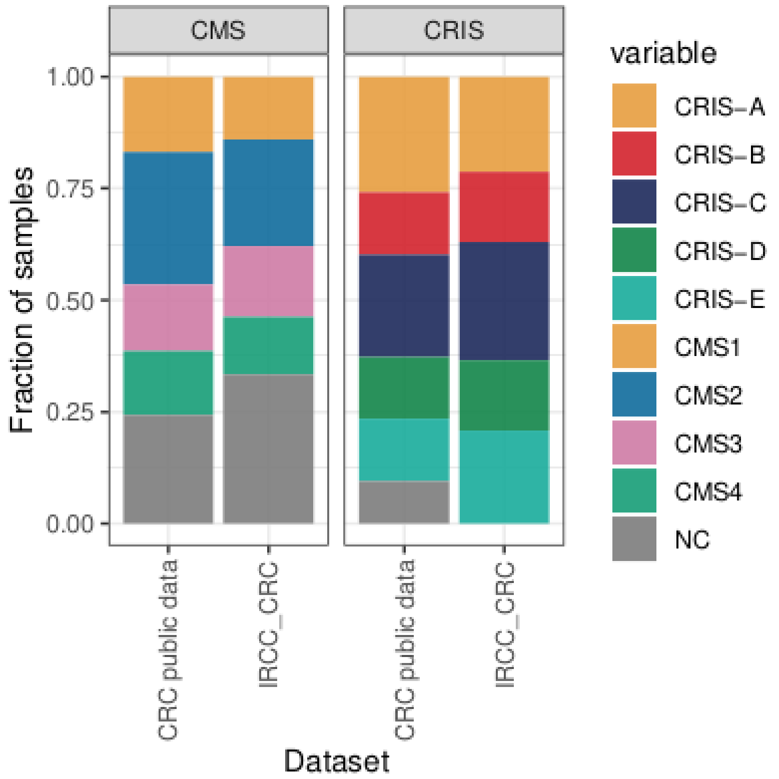


***Figure S9 - Comparing CMS/CRIS expression class frequency in IRCC-PDX and TCGA COAD/READ***

*CMS and CRIS transcriptional class frequencies observed in publicly available CRC RNAseq profiles of tumours (see below) or our full IRCC-PDX (N=442).*

*CRIS classes were obtained from [Isella 2017 https://doi.org/10.1038/ncomms15107] for the following datasets: TCGA, GSE13067, GSE13294, GSE14333, GSE17536, GSE20916,GSE2109, GSE23878, GSE33113, GSE35896, GSE37892, GSE39582, GSE5851, GSE59857, KFSYSCC, PETACC (https://pubmed.ncbi.nlm.nih.gov/33001764/). CMS classes were obtained from [Eide 2017 https://doi.org/10.1038/s41598-017-16747-x] for the following datasets: TCGA, Medico et al. (2015) (cell lines), van de Wetering et al. (2015) (organoids), Fujii et al. (2016) (organoids), Julien et al. (2012) (PDX), Uronis et al. (2012) (PDX), Gao et al. (2015) (PDX), Isella et al. (2017) (PDX).*

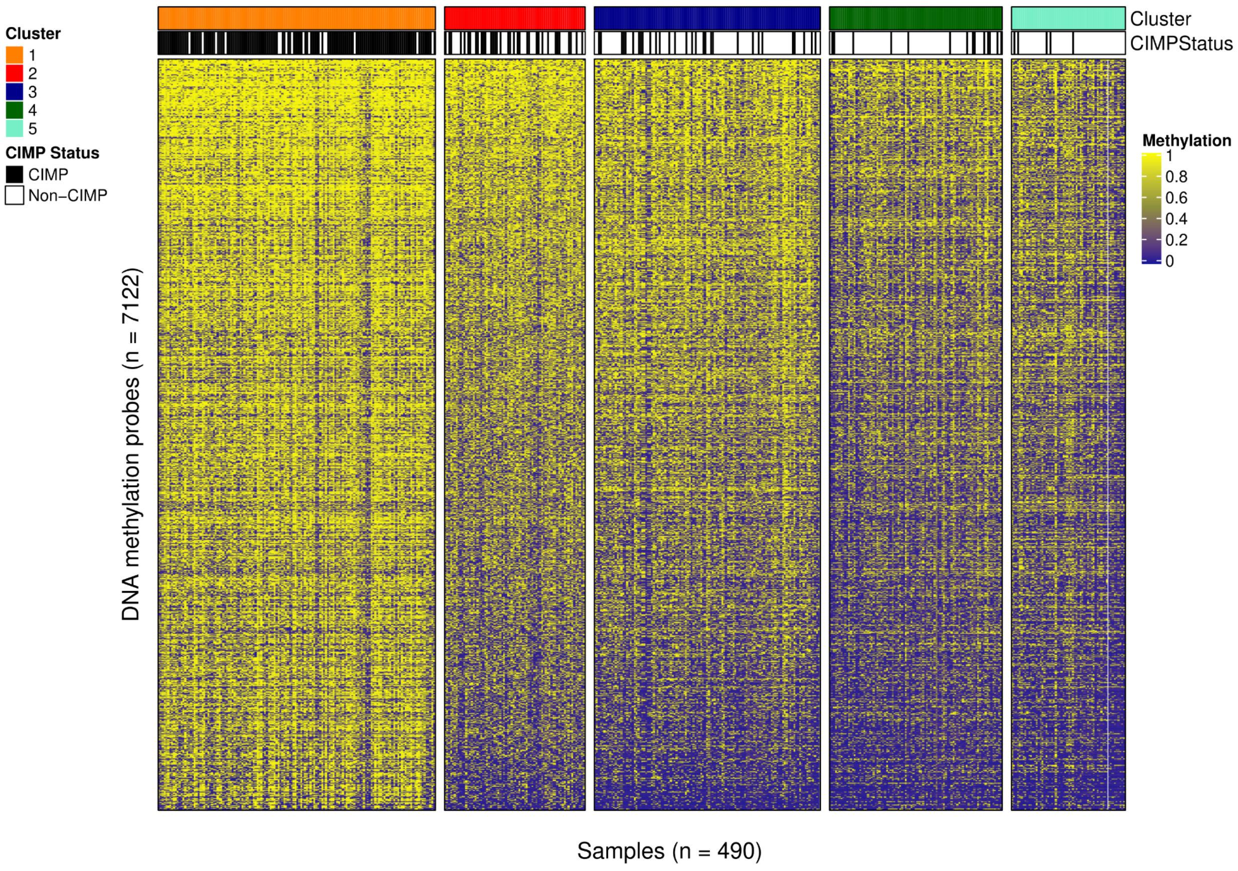


***Figure S10 - Probes methylation status in PDXs.*** *Methylation levels of 7,122 most variant probes (on the row) in the IRCC-PDXs cohort (490 samples with available methylation data on the columns), (see Fig.S6 for CIMP annotation). Samples are grouped according to non negative matrix derived clusters (as indicated by the different colours). The first cluster is significantly more hypermethylated over all measured CpG islands (median beta methylation level = 0.81, Kruskal-Wallis test p-value < 2.2e-16).*

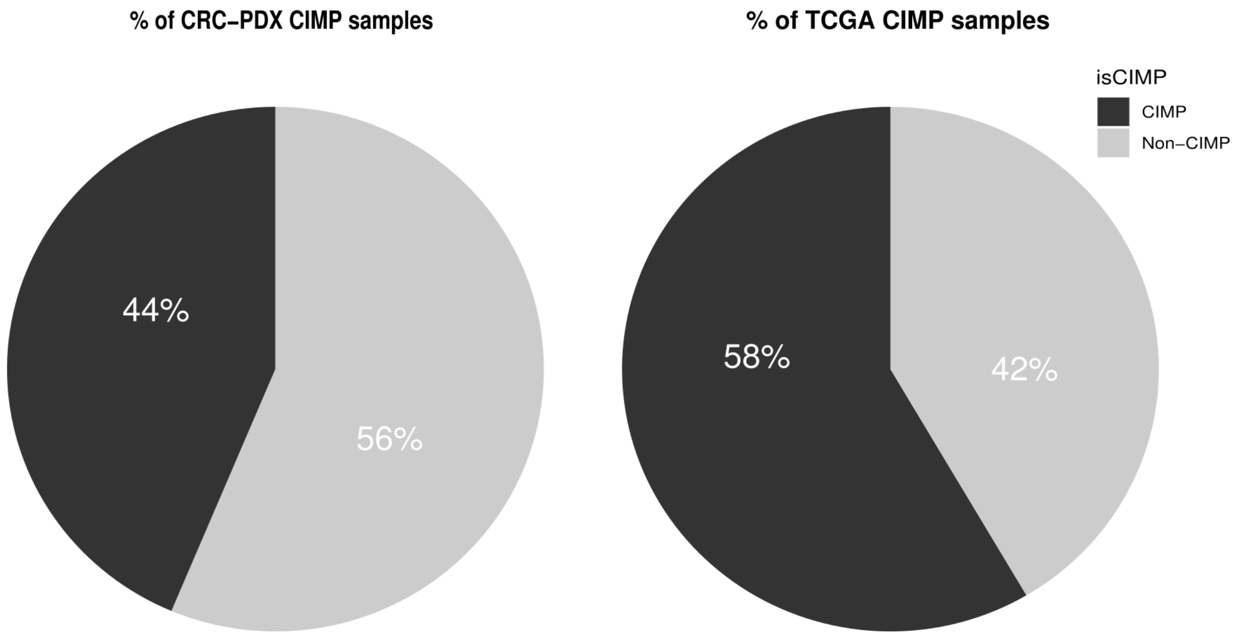


***Figure S11 - CIMP samples in IRCC-PDX and TCGA*** *CpG island methylator phenotype (CIMP) and Non-CIMP class frequency in our IRCC-PDX cohort and TCGA COAD/READ obtained using Hinoue et al., 2012 (http://www.genome.org/cgi/doi/10.1101/gr.117523.110.) markers panel. This panel consists of B3GAT2, FOXL2, KCNK13, RAB31, and SLIT1 genes. Here, DNA methylation of three or more of these markers qualifies a sample as CIMP, with a  -value threshold of ≥ 0.1. We are able to identify consistent percentages among the two datasets: 214 CIMP samples (44%) in our cohort, and 235 (58%) in TCGA.*

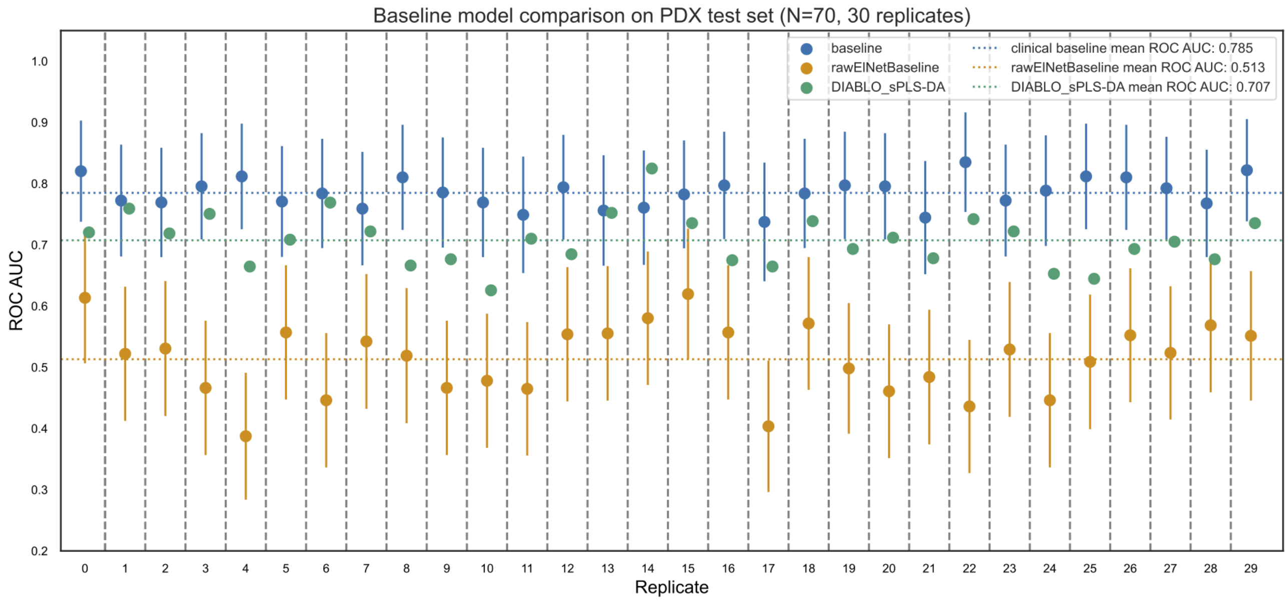


***Figure S12 - The clinical signature baseline is the stricter benchmark for our stacked classifier****. This figure compares the performance of three baseline models over 30 holdout shuffle replicates of our IRCC-PDX dataset, using ROC AUC confidence intervals (vertical bars) computed using DeLong's method. The clinical signature elastic net classifier ("baseline", in blue) is based on 4 binary features: the mutational status of KRAS,BRAF,NRAS, and the right-colon origin of the primary tumour. The DIABLO sPLS-DA classifier ("DIABLO_sPLS-DA", in green) is implemented in MixOmics and only returns ROC AUC point estimates (see Methods). The raw elastic net baseline ("rawElNetBaseline", in orange) uses the full set of raw genomic, transcriptomic, clinical, and the NMF clustered methylonics features as input (39960 features in total). We observe that the clinical baseline outperforms DIABLO sPLS-DA by 0.08 ROC AUC on average and the raw elastic net baseline by 0.27 ROC AUC on average.*

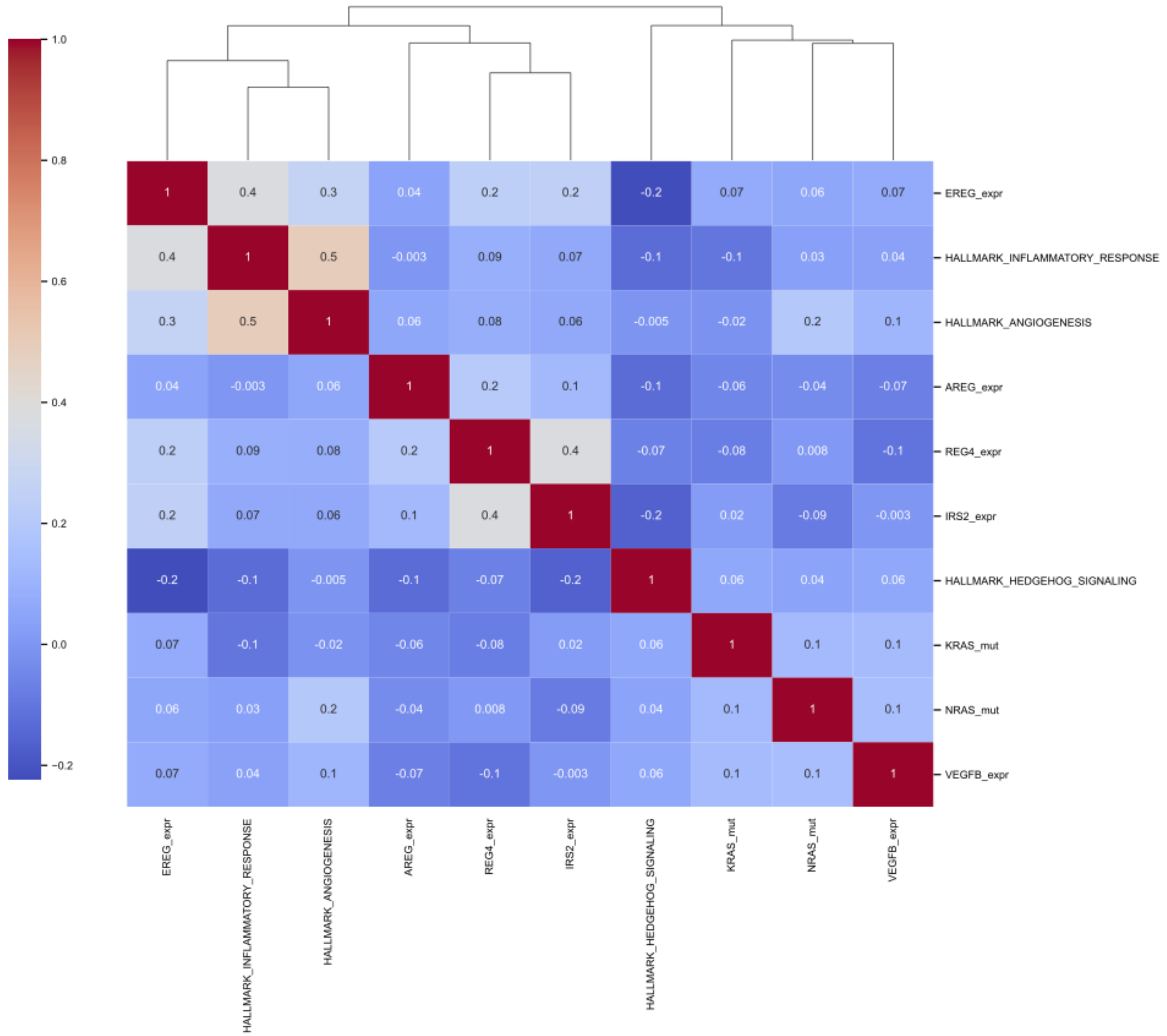


***Figure S13 - Feature value correlation in the IRCC-PDX dataset**. This heatmap shows the relationship among top feature (as per Fig 4a) values in the IRCC-PDX dataset (train set, N=231) as measured by Perarson correlation. We detect no collinearity (Pearson coefficient > 0.7) among these features. The strongest correlation (0.5) is observed between the "inflammatory response" and "angiogenesis" ssGSEA gene set scores. The highest anticorrelation (-0.2) is observed between the "hedgehog signalling: ssGSEA score and EREG's expression.*

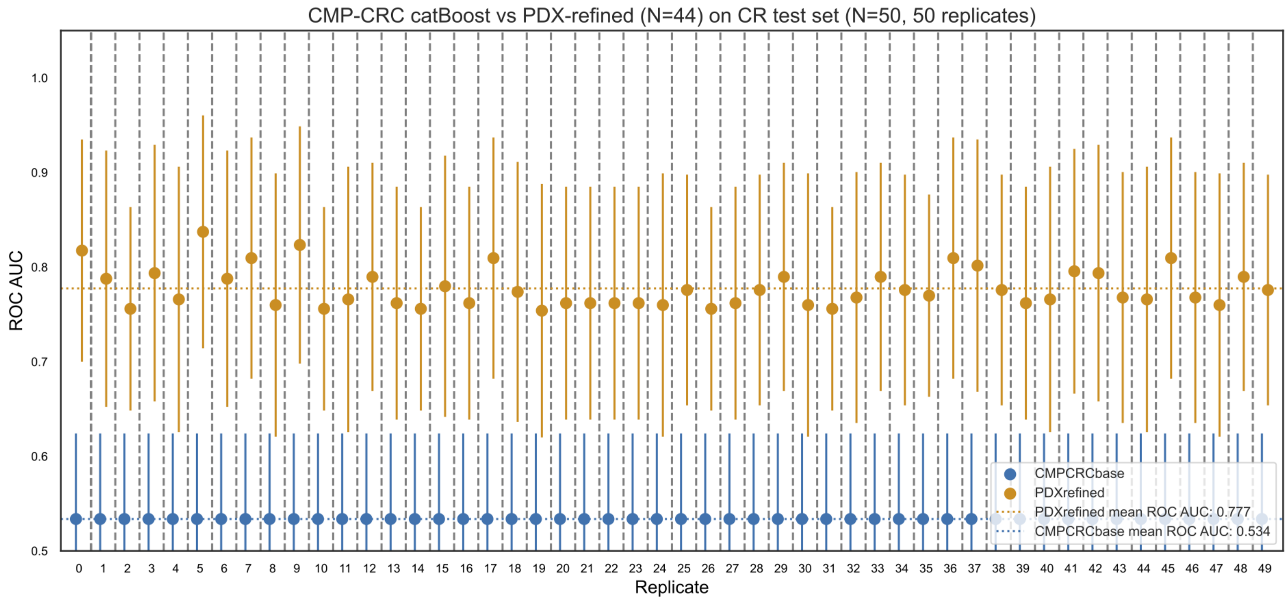


***Figure S14 - Classifiers refined on CRC PDXs outperform those trained on CRC 2d cell lines only****.*
*This figure compares ROC AUC confidence intervals (vertical bars, obtained via DeLong's method) across 50 replicates where a CatBoost classifier of Cetuximab sensitivity is first trained on all 44 CRC cell lines available in the CMP dataset, and then tested on the CR-PDX dataset ('CMPCRCbase', in blue). We then continue training ('refine') this same 'CMPCRCbase' model on a downsampled set of 44 PDXs from our IRCC -DX dataset, followed by another test on the CR-PDX dataset ('PDXrefined', in orange). Here, we observe a noticeable improvement in predictive performance after the cell line trained classifiers are refined on a set of PDX examples of equal size (N=44), to the point that their ROC AUC confidence intervals do not overlap.*

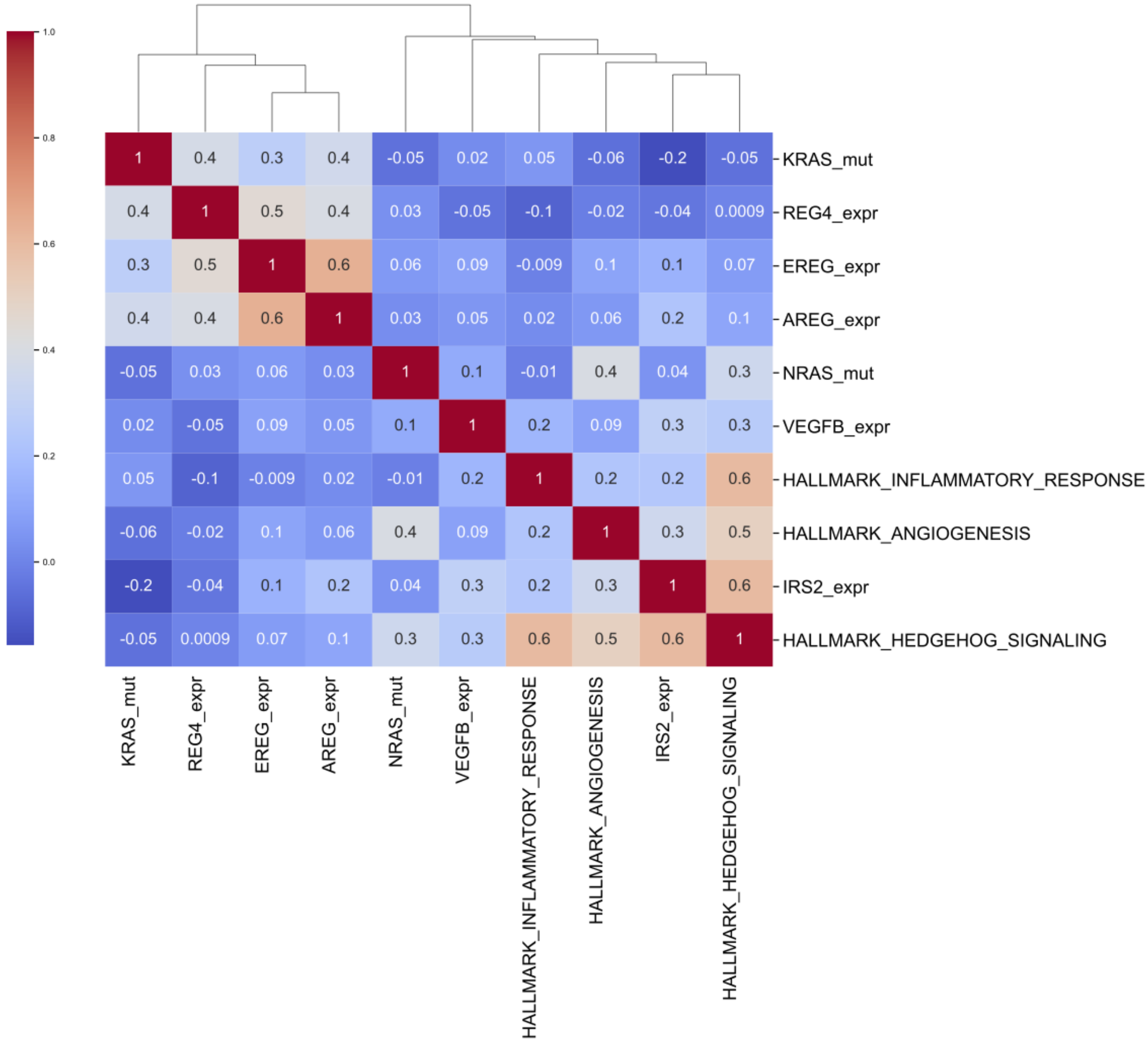


***Figure S15 - CeSta SHAP value correlation in the CR-PDX dataset****. This heatmap shows the relationship among CeSta classifier SHAP values (Methods) computed using the IRCC-PDX dataset (train set, N=231) for a selection of top features, as measured by Pearson correlation. Features whose SHAP values are highly correlated influence the classifier's output in the same direction.*

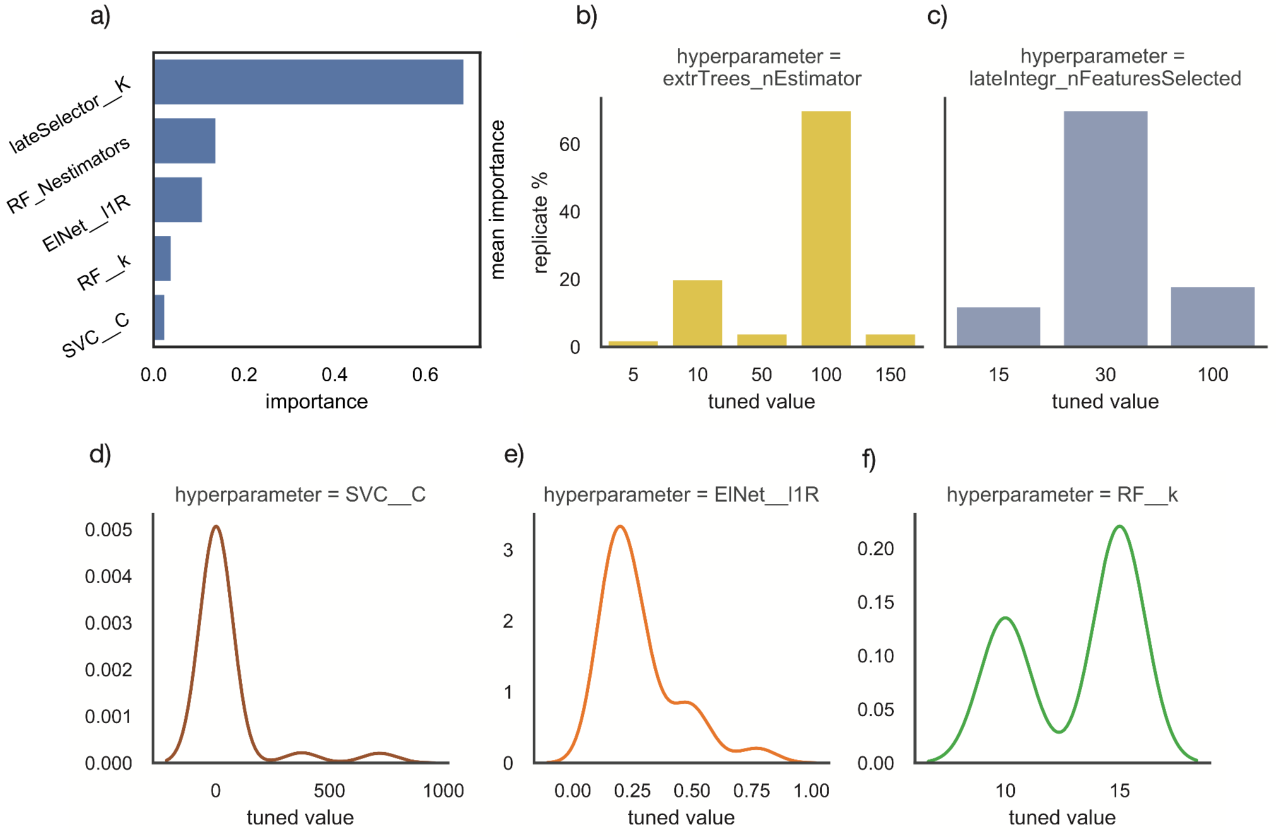


***Figure S16 - CeSta hyperparameter importance and tuned value distribution across 50 independently tuned replicates. a)*** *Shows mean hyperparameter importance (as computed by Optuna) across 50 independent tuning runs (internal validation) for the top 5 CeSta hyperparameters.* ***b,c)*** *Illustrate tuned value frequency for two categorical hyperparameters: the number of estimators in our extraTrees lvl 1 classifier, and the number of top features selected by our first late-integration filter, respectively (Methods,* ***Fig 1b****), across 50 independent tuning runs.* ***d-f)*** *Report tuned value density for three continuous hyperparameters: regularisation parameter of our SVC, Elastic-Net mixing parameter for our Elastic-Net penalised logistic classifier, and number of features selected by our extraTrees classifier pipeline filter, respectively, across 50 independent tuning runs.*

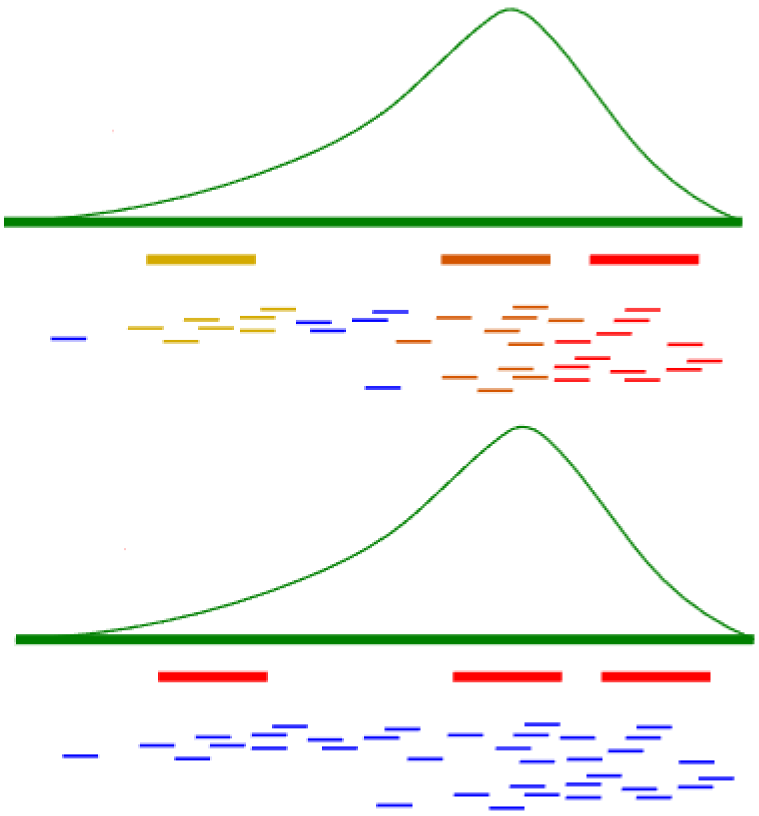

***Figure S6 - Mapping copy number segments to GISTIC/ADMIRE events and intOGen genes.*** *This diagram illustrates our process for mapping TCGA COAD/READ copy number segments (in blue) to a GISTIC/ADMIRE copy number variation event (in green) and to intOGen genes (in red, orange, and yellow) using BEDtools (Methods).*

| | FE_oddsR | FE_Pval | ate | percent_lift | support | abs_percent_lift | FE_adj_Pval | FE_FDR<.05 | FE_adj_FDR<.05 | U1 | U1_Pval | U1_adj_Pval | U1_FDR<.05 | U1_adj_FDR<.05 | singleOmic_logit_Pval | singleOmic_logit_coeff | omic |
|---|---|---|---|---|---|---|---|---|---|---|---|---|---|---|---|---|---|
| **cluster_1** | 0.163 | 0.000 | -0.273 | -0.768 | 34.000 | 0.768 | 0.000 | True | True | | | | | | 0.001 | -1.558 | Meth |
| **cluster_4** | 3.600 | 0.002 | 0.221 | 1.682 | 40.000 | 1.682 | 0.022 | True | True | | | | | | 0.036 | 0.891 | Meth |
| **KRAS_mut** | 0.174 | 0.000 | -0.331 | -0.719 | 46.000 | 0.719 | 0.000 | True | True | | | | | | 0.082 | -1.360 | Gen |
| **BRAF_mut** | 0.079 | 0.003 | -0.120 | -0.911 | 11.000 | 0.911 | 0.042 | True | True | | | | | | 0.065 | -2.059 | Gen |
| **NRAS_mut** | 0.139 | 0.053 | -0.067 | -0.851 | 7.000 | 0.851 | 0.686 | False | False | | | | | | 0.069 | -2.043 | Gen |
| **CLSPN_mut** | 0.101 | 0.014 | -0.093 | -0.888 | 9.000 | 0.888 | 0.176 | True | False | | | | | | 0.021 | -3.042 | Gen |
| **FGFR1_highGain_cnv** | 5.538 | 0.001 | 0.183 | 3.471 | 24.000 | 3.471 | 0.019 | True | True | | | | | | 0.015 | 1.723 | Gen |
| **mut_group_7** | 0.101 | 0.014 | -0.093 | -0.888 | 9.000 | 0.888 | 0.176 | True | False | | | | | | 0.482 | -0.930 | Gen |
| **mut_group_16** | 0.117 | 0.027 | -0.080 | -0.872 | 8.000 | 0.872 | 0.350 | True | False | | | | | | 0.539 | -0.829 | Gen |
| **mut_group_5** | 0.336 | 0.033 | -0.114 | -0.617 | 20.000 | 0.617 | 0.433 | True | False | | | | | | 0.764 | 0.277 | Gen |
| **mut_group_12** | 7.792 | 0.036 | 0.081 | 6.153 | 9.000 | 6.153 | 0.474 | True | False | | | | | | 0.088 | 2.287 | Gen |
| **IRINOTECAN-based treatments (Y/N) (includes FOLFIRI, XELIRI +/- other drugs)** | 1.821 | 0.260 | 0.071 | 0.676 | 23.000 | 0.676 | 3.383 | False | False | | | | | | 0.224 | 0.576 | Clin |
| **Right** | 0.516 | 0.125 | -0.112 | -0.404 | 35.000 | 0.404 | 1.626 | False | False | | | | | | 0.098 | -0.646 | Clin |
| **HALLMARK_UV_RESPONSE_DN** | | | -0.084 | -6.637 | 80.000 | 6.637 | | | | 2,402.500 | 0.005 | 0.082 | True | False | 0.241 | -1.743 | Expr |
| **HALLMARK_KRAS_SIGNALING_DN** | | | 0.011 | -2.241 | 84.000 | 2.241 | | | | 3,330.500 | 0.735 | 11.758 | False | False | 0.096 | 3.995 | Expr |
| **HALLMARK_INFLAMMATORY_RESPONSE** | | | -0.095 | -3.771 | 68.000 | 3.771 | | | | 2,348.500 | 0.003 | 0.046 | True | True | 0.228 | 2.246 | Expr |
| **HALLMARK_EPITHELIAL_MESENCHYMAL_TRANSITION** | | | -0.060 | -4.102 | 74.000 | 4.102 | | | | 2,587.500 | 0.030 | 0.475 | True | False | 0.837 | -0.339 | Expr |
| **HALLMARK_APICAL_JUNCTION** | | | -0.036 | 11.157 | 63.000 | 11.157 | | | | 2,867.500 | 0.220 | 3.524 | False | False | 0.925 | 0.202 | Expr |
| **HALLMARK_PANCREAS_BETA_CELLS** | | | -0.049 | -1.491 | 83.000 | 1.491 | | | | 2,860.500 | 0.211 | 3.384 | False | False | 0.979 | -0.025 | Expr |
| **HALLMARK_WNT_BETA_CATENIN_SIGNALING** | | | -0.021 | 0.684 | 65.000 | 0.684 | | | | 3,011.500 | 0.460 | 7.366 | False | False | 0.704 | -0.419 | Expr |
| **HALLMARK_ANGIOGENESIS** | | | -0.095 | -1.992 | 80.000 | 1.992 | | | | 2,329.500 | 0.002 | 0.037 | True | True | 0.291 | -1.455 | Expr |
| **HALLMARK_MYOGENESIS** | | | -0.013 | 15.020 | 76.000 | 15.020 | | | | 3,068.500 | 0.586 | 9.370 | False | False | 0.428 | -1.952 | Expr |
| **HALLMARK_HEDGEHOG_SIGNALING** | | | -0.085 | -5.444 | 69.000 | 5.444 | | | | 2,426.500 | 0.007 | 0.105 | True | False | 0.705 | -0.507 | Expr |
| **HALLMARK_XENOBIOTIC_METABOLISM** | | | -0.001 | 0.018 | 63.000 | 0.018 | | | | 3,353.500 | 0.677 | 10.833 | False | False | 0.181 | 1.727 | Expr |
| **PROGENy_EGFR** | | | 0.294 | -1.870 | 102.000 | 1.870 | | | | 3,892.500 | 0.025 | 0.400 | True | False | 0.188 | 0.342 | Expr |
| **PROGENy_MAPK** | | | -0.025 | -1.542 | 24.000 | 1.542 | | | | 3,610.500 | 0.198 | 3.171 | False | False | 0.716 | 0.087 | Expr |
| **REG4_expr** | | | -1.691 | -0.190 | 157.000 | 0.190 | | | | 2,111.500 | 0.000 | 0.002 | True | True | 0.001 | -0.369 | Expr |
| **EREG_expr** | | | 0.921 | 0.099 | 157.000 | 0.099 | | | | 4,488.500 | 0.000 | 0.000 | True | True | 0.001 | 0.313 | Expr |
| **mutational_burden** | | | -1.930 | -0.225 | 161.000 | 0.225 | | | | 2,481.500 | 0.011 | 0.172 | True | False | 0.506 | -0.047 | Gen |

***Table S1 - Association between selected features and the Cetuximab sensitivity****. This table illustrates multiple statistical metrics of association between 30 features selected from the corresponding training set in the first step of our stacked classifier pipeline and the target variable. Specifically "FE_oddsR" and "FE_Pval" correspond to Fisher's exact test odds ratios and p-values computed against the target labels for binary features. "U1" and "U1_Pval" correspond to Mann-Whitney U test statistics and p-values computed against the target for continuous features. "ATE" (average treatment effect) corresponds to the difference of the mean value of each feature among Cetuximab responders and non responders. "Percent_lift" is the ATE divided by the mean value of the corresponding feature among Cetuximab non responders and "abs_percent_lift" to the absolute value of the percent lift for this feature. "Support" counts the number of PDXs in which the feature is either observed (if binary) or non null (if continuous). "singleOmic_logit_Pval" and "singleOmic_logit_coeff" correspond to the p-values and coefficients of a logit model fit independently to each omic feature set ("omic")*
